# Endothelial transmigration hotspots limit vascular leakage through heterogeneous expression of ICAM1

**DOI:** 10.1101/2022.01.14.476297

**Authors:** Max L.B. Grönloh, Janine J.G. Arts, Sebastián Palacios Martínez, Amerens A. van der Veen, Lanette Kempers, Abraham C.I. van Steen, Joris J.T.H. Roelofs, Martijn A. Nolte, Joachim Goedhart, Jaap D. van Buul

## Abstract

Upon inflammation, leukocytes leave the circulation by crossing the endothelial monolayer at specific transmigration ‘hotspot’ regions. Although these regions support leukocyte transmigration, their functionality is not clear. We found that endothelial hotspots function to limit vascular leakage during transmigration events. Using the photo-convertible probe mEos4b, we traced back and identified original endothelial transmigration hotspots. Using this method, we show that the heterogeneous distribution of ICAM-1 determines the location of the transmigration hotspot. Interestingly, loss of ICAM-1 heterogeneity either by CRISPR/Cas9-induced knockout of ICAM-1 or equalizing the distribution of ICAM-1 in all endothelial cells results in loss of TEM hotspots but not necessarily in reduced TEM events. Functionally, loss of endothelial hotspots results in increased vascular leakage during TEM. Mechanistically, we demonstrate that the 3 extracellular Ig-like domains of ICAM-1 are crucial for hotspot recognition. However, the intracellular tail of ICAM-1 and the 4^th^ Ig-like dimerization domain are not involved, indicating that intracellular signalling or ICAM-1 dimerization is not required for hotspot recognition. Together, we discovered that hotspots function to limit vascular leakage during inflammation-induced extravasation.

## Introduction

The migration of leukocytes towards sites of infection or tissue damage is key to the inflammatory response of the innate immunity. To reach the underlying tissue, leukocytes exit the circulation through a process called transendothelial migration (TEM). TEM consists of several subsequent steps, known as the multistep process, introduced by Butcher and Springer, and although the basis of this concept is still very solid, new details still are discovered and characterized, adding to the full picture of the multistep paradigm^1–6^.

It is recognized that during inflammation, when leukocytes extravasate, vessels do not leak^2, 7^. Work by our group and others showed that the gaps induced by the penetrating leukocytes are quickly repaired by a variety of intracellular processes within the endothelium itself^8–10^. Intravital microscopy revealed that neutrophil migration through the endothelial monolayer and the basement membrane and pericyte sheath does not occur randomly, but in fact occurs at predefined exit sites, called “hotspots”^11^. Although it is without a doubt that leukocytes use hotspots to cross the endothelium and many factors have been proposed to determine hotspot composition and localization^12^, the physiological relevance why leukocytes would prefer to cross the endothelium at hotspots is unclear. Examples of hotspot regulators are heterogenous chemokine gradients^13, 14^, differences in substrate stiffness^15, 16^, junction phenotype^17^ and recently reported varying junctional membrane protrusion activities between individual endothelial cells^18^ and autophagy at junction regions^19^. Additionally, the composition and density of the pericyte sheath and the basement membrane layer may also influence the location of both endothelial and basement membrane hotspots^20, 21^.

As endothelial adhesion molecules are important regulators of efficient neutrophil TEM, this protein family may also play a role in the localization of TEM hotspots. Intercellular adhesion molecule (ICAM)-1 and ICAM-2, both heavily involved in neutrophil adhesion, are transmembrane glycoproteins of the immunoglobulin superfamily and in particular ICAM-1 is highly upregulated on inflamed endothelium^22^. ICAM-1 has several splicing variants, but generally consists of 5 extracellular immunoglobulin (Ig)-like domains, the fourth one regulating homodimerization^23, 24^. ICAM-2 has just two Ig-like domains, which are homologues to the first and second Ig-like domains of ICAM-1^25^. Both ICAM-1 and ICAM-2 bind neutrophil integrins lymphocyte function associated antigen 1 (LFA-1) (CD11a/CD18) and macrophage-1 antigen (Mac-1) (CD11b/CD18) and are mainly involved in the firm adhesion and crawling steps of the endothelium^26, 27^. LFA-1 has been reported to bind to the first extracellular domains of ICAM-1 and ICAM-2^28, 29^. Mac1 binds the third extracellular domain of ICAM-1, whereas information on association between Mac1 and ICAM-2 is scarce^30^. Both ICAM-1 and -2 are linked to the actin cytoskeleton via modulators such as α-actinin-4, filamin B and cortactin^31, 32^. After inflammation, ICAM-1 is upregulated and displays a typical patchy pattern *in vitro* as well as *in vivo*^33–35^. In contrast to ICAM-1, ICAM-2 is already constitutively expressed on the endothelium in normal conditions^36^.

The biological relevance of TEM hotspots is not yet understood. It has been hypothesized that the interaction of leukocytes with only a small selection of the vessel could help maintain the barrier integrity of the endothelium^11^, but no evidence in favour of this theory has been put forward. It has been shown that transmigration of neutrophils is not correlated with local leakage at those specific sites, as mechanisms exist to limit transmigration-induced vascular leakage^9, 10^. Other studies have shown that leukocyte adhesion to the endothelium itself can already trigger vascular leakage^37^, suggesting that adhesion-related processes, and not diapedesis-related ones, can be associated with vascular leakage.

Here, we reveal the biological relevance of neutrophil TEM hotspots in the endothelial monolayer by establishing a molecular link between endothelial hotspots that are regulated by ICAM-1 distribution and vascular leakage. Mechanistically, we show that ICAM-1 heterogeneity is crucial for the presence of TEM hotspots. Loss of ICAM-1 heterogeneity either by CRISPR/Cas9 knockout or distributing and expressing ICAM-1 at equal levels in all endothelial cells results in loss of TEM hotspots but not in altered neutrophil adhesion or diapedesis efficacy. Interestingly, we found under these conditions an increase in vascular leakage during TEM. Hotspot functionality and recognition depends on the first 3 extracellular Ig-like domains of ICAM-1 but not on its intracellular tail or the 4^th^ Ig-like dimerization domain. This indicates that intracellular signalling or ICAM-1 dimerization is not required for hotspot functionality and recognition. Restoration of the heterogeneous distribution of ICAM-1 rescued the increase in local permeability during diapedesis.

Thus, our study reveals the functional importance of endothelial heterogeneity of adhesion molecules in regulating TEM hotspots under inflammatory conditions and these hotspots function to limit vascular leakage during leukocyte extravasation.

## Results

### Transmigration hotspots exist *in vitro*

*In vivo* data show that leukocytes prefer local exit sites, named transendothelial migration (TEM) hotspots, although the functionality of the hotspots is not clear^11^. To investigate the functionality of hotspots, we first need to confirm that these hotspots also exist *in vitro.* Neutrophil transmigration was studied under physiological flow conditions using tumor necrosis factor (TNF)-α-stimulated human umbilical vein endothelial cells (HUVEC). We observed that neutrophils prefer to leave the endothelium at specific sites, while almost completely ignoring other areas (Figure 1A). To quantify if more than one neutrophil preferred the same endothelial area to transmigrate, we used an unbiased ‘nearest neighbour’ analysis, a method very suitable to detect clustering of spatial data. We calculate the mean distance of each transmigration event to three transmigration events that were nearest and compared this to a same number of randomly generated spots (Figure 1B). Indeed, a significant decrease in the average distance to the 3 nearest transmigration events compared to the randomized spots was found, indicative for the existence of preferred endothelial hotspots that regulate transmigration events (Figure 1C & S1A). To check if the number of neighbours used in the analysis did not influence the outcome, we also analysed transmigration events that were closest to one, five and nine nearest transmigration events and found similar patterns (Figure S1B). Thus, from these quantitative analyses, we conclude that TEM hotspots also exist *in vitro*.

**Figure 1.**
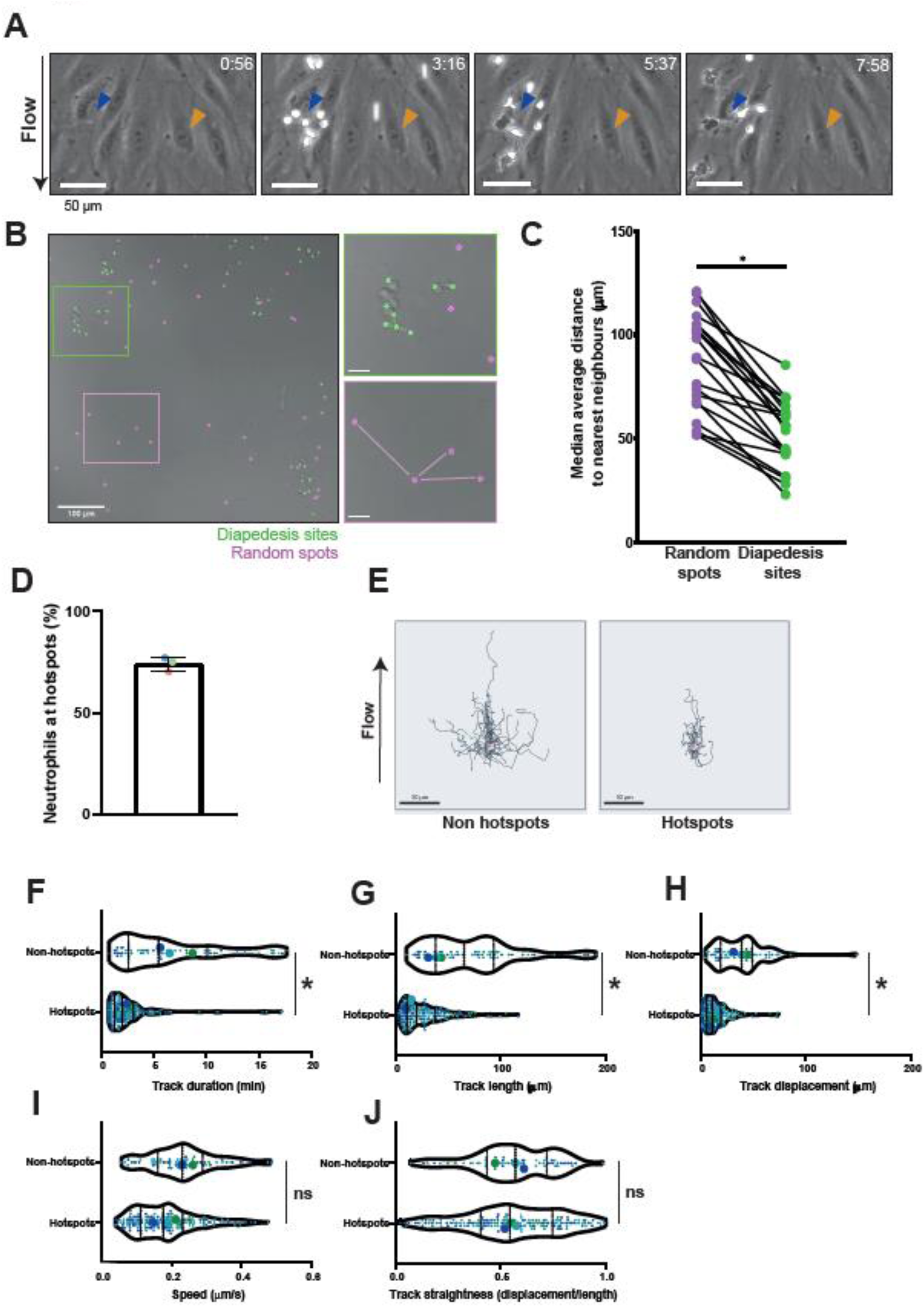
*Neutrophils transmigrate more efficient at TEM hotspots*. **(A)** Still from a time-lapse TEM assay, showing a neutrophil TEM hotspot indicated with a blue arrow and a largely by neutrophils ignored region indicated with an orange arrow. The direction of flow is from top to bottom, time is indicated in minutes at the top right. Scale bar, 50 µm. **(B)** Brightfield still image from a neutrophil TEM experiment, with marked the diapedesis sites (green) and computationally generated random spots (magenta). Scale bar, 100 µm **(C)** Medians of average distance to 3 nearest neighbours for each timelapse are plotted. Medians are paired with medians of average distance to 3 nearest neighbours of corresponding randomly generated spots. Paired t-test on the 21 medians: p<0.0001. **(D)** Bar graph of total percentage of neutrophils utilizing hotspots (≥2 diapedesis events). Means from 3 independent experiments are shown. Bar graph shows mean with SD. **(E)** 40 overlayed tracks of crawling neutrophils that eventually transmigrate at a TEM hotspot (right, ≥2 diapedesis events) or not (left, 1 diapedesis event). Scale bar, 50 µm**. (F,G,H,I,J)**. Small dots represent individual datapoints, large dots are medians from each experiment. 174 hotspot tracks and 62 non-hotspot tracks from 3 independent experiments are represented in 3 different colours. Paired t-test on the medians of 3 independent experiments. **(F)** Violin plot of track duration of crawling neutrophils that eventually transmigrate at a TEM hotspot or not. p=0.0121. **(G)** Violin plot of total track length of crawling neutrophils that eventually transmigrate at a hotspot or not. p = 0.039. **(H)** Violin plot of the total displacement (distance between the begin and the end of the track) of crawling neutrophils that eventually transmigrate at a TEM hotspot or not. p = 0.019. **(I)** Violin plot of average speed of crawling neutrophils that eventually transmigrate at a TEM hotspot or not. p = 0.051. **(J)** Violin plot of Track straightness (displacement from (F)/length from (E)) of crawling neutrophils that eventually transmigrate at a TEM hotspot or not. p = 0.95.

To study whether neutrophils at TEM hotspots displayed different spatiotemporal crawling dynamics compared to neutrophils that ignored hotspots, we classified neutrophil crawling tracks in physiological flow time-lapse recordings and classified a ‘hotspot track’ when a neutrophil would transmigrate within 50 µm, the average diameter of one endothelial cell of another neutrophil (Figure S1C). We found that around 75% of all TEM events occurred at a hotspot location (Figure 1D). Moreover, neutrophils that used TEM hotspots crawled for shorter distances and shorter durations than neutrophils that did not undergo diapedesis at hotspots, although migration speed and crawling linearity were not altered (Figure 1E-J). Combined, these data indicate that TEM occurs more frequently and is more efficient at endothelial hotspots.

### ICAM-1 marks neutrophil TEM hotspots

To understand how neutrophils find these hotspots, we need to be able to identify endothelial TEM hotspots once neutrophils have used them. To do so, endothelial cells (ECs) were transfected with the photoconvertible probe mEos4b and neutrophil TEM under flow was monitored in real time. Sites of TEM were determined on the fly and marked as field of view (FOV). FOV were exposed to 405 nm light, converting mEos4b to a red fluorescent protein. The red fluorescence allowed us to trace back and identify original TEM hotspots to screen for candidate adhesion molecules (Figure 2A). Using this technique, we found that the distribution of ICAM-1 perfectly correlated with TEM hotspots, whereas VCAM-1 and ICAM-2 did not (Figure 2B).

**Figure 2.**
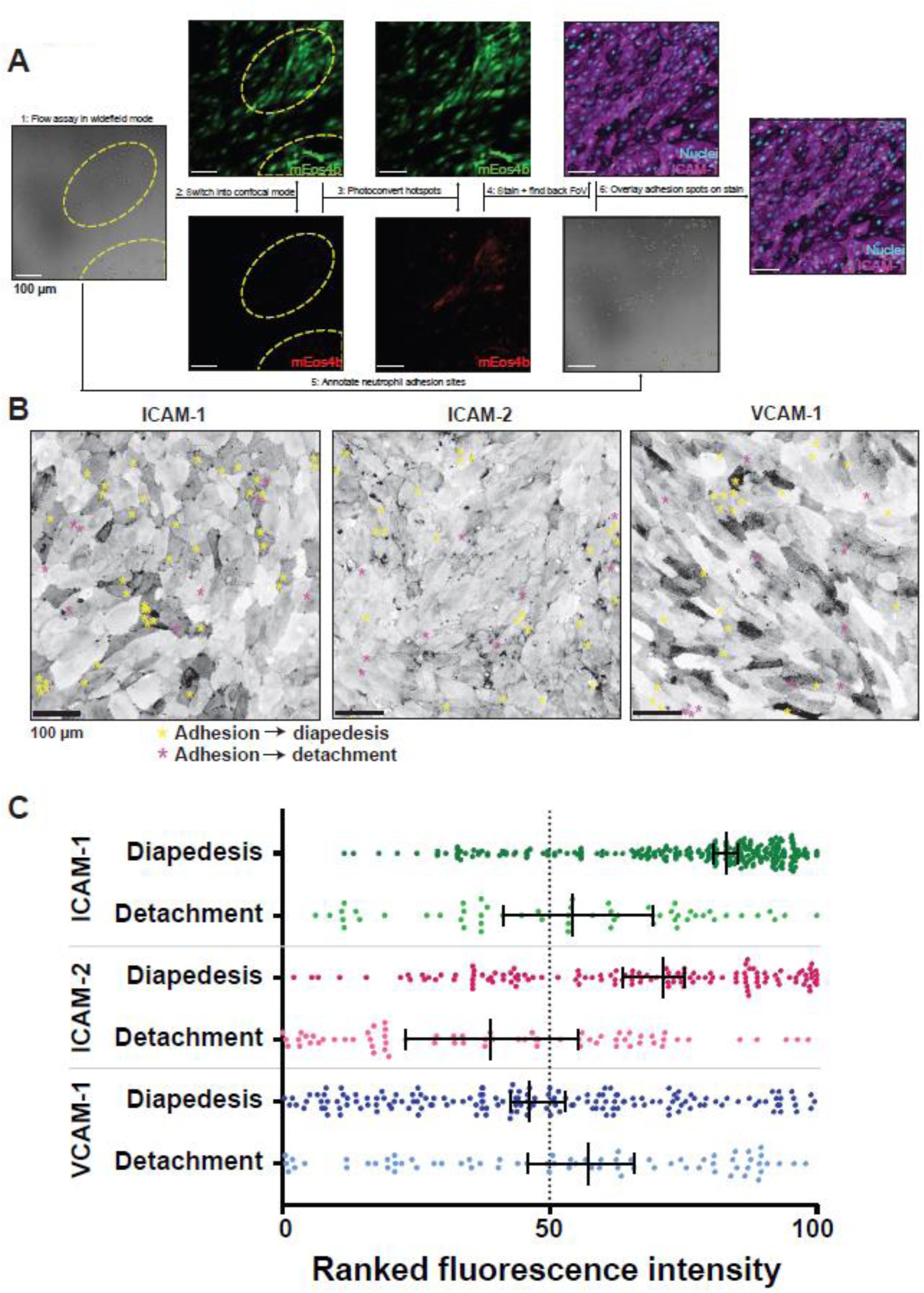
*Neutrophil transmigration hotspots are located at ICAM-1 high expressing cells.* **(A)** Simplified workflow for photoconversion experiments. (1) neutrophil flow assays were performed in widefield mode with HUVECs expressing mEos4b. (2) The same field of views were imaged in confocal mode. (3) Areas where hotspots had appeared in the widefield video were converted from green to red using a 405 nm laser. (4) The slides were fixed and the nuclei and an adhesion molecule were stained. The original fields of view were found backing by looking for red signal. (5) In the original video, all successful and unsuccessful adhesion events were annotated. (6) The adhesion spots were overlayed with the stained image. **(B)** Inverted greyscale LUT of immunofluorescence stains on HUVECs for ICAM-1, ICAM-2 and VCAM-1 of areas of which time-lapses were made, relocated by photoconversion of mEos4b. With yellow asterisks, adhesion events that led to diapedesis are shown. With magenta asterisks, adhesion events that were followed by detachment are shown. Scale bar = 100 μm. **(C)** Quantification of adhesion events on HUVEC cells ranked by their preference for diapedesis for expression levels of ICAM-1 (green dots, n = 254 diapedesis adhesion events and n = 57 detachment adhesion events), ICAM-2 (magenta dots n = 129 diapedesis adhesion events and n = 79 detachment adhesion events) and VCAM-1 (blue dots n = 163 diapedesis adhesion events and n = 72 detachment adhesion events). Data shown is from 9 (ICAM-1 and VCAM-1) or 6 (ICAM-2) images from 3 independent experiments. Median with 95% CI is shown.

To validate these observations, we identified individual ECs, stained them for ICAM-1, -2 or VCAM-1 and ranked them by fluorescence intensity, representing surface protein expression levels. Next, we correlated TEM sites to fluorescence intensity, plotted all TEM events, and discriminated between neutrophils that transmigrated (marked with yellow asterisks) and neutrophils that did adhere to the endothelium, but detached again (marked with magenta asterisks) (Figure 2A). These data showed that most adhesion events that led to successful TEM required ICAM-1^high^ ECs (Figure 2C). Interestingly, we also found a preference for ICAM-2^high^ ECs, albeit less prominent (Figure 2C), suggesting that neutrophils can differentiate between high and low ICAM-2-expressing ECs. Neutrophil adherence to VCAM-1 was completely random, underscoring the fact that neutrophils did not express VCAM-1-counter receptor VLA-4 (Figure 2C). These unbiased data indicate that ICAM-1 marks TEM hotspots.

### Heterogeneous distribution of endothelial adhesion molecules

Based on the strong correlation between ICAM-1^high^ expression and TEM hotspots, we hypothesize that the heterogeneous expression of endothelial ICAM-1 leads to increased neutrophil adhesion and thus TEM at those sites. To examine adhesion molecule distribution within an inflamed endothelial monolayer in more detail, we stained TNF-α-treated ECs for ICAM-1, ICAM-2 and VCAM-1 and found that ICAM-1 expression is distributed in a heterogenous manner: some ECs displayed high levels of ICAM-1, whereas others did not (Figure 3A). Furthermore, ICAM-1 localized to apical filopodia ^38, 39^, but this was only observed in ICAM-1^high^ ECs. In contrast to ICAM-1, ICAM-2 showed a much more homogenous distribution within the endothelial monolayer and was slightly enriched at junction areas but not at filopodia (Figure 3A). Finally, VCAM-1 did show heterogenous expression and was enriched in filopodia on ECs that showed VCAM-1^high^ expression (Figure 3A). To quantify heterogeneous distribution, we measured fluorescence intensity of individual ECs and normalized the fluorescent values within each field of view to correct for variation between images. Indeed, ICAM-1 and VCAM-1 showed a wide distribution of the violin plot, indicating increased heterogeneous distribution, with nuclei staining as maximal equal distribution (Figure 3B). This quantification allowed us to measure the variation of protein distribution within one EC monolayer. ICAM-1 and VCAM-1 showed strong heterogenous distribution, whereas ICAM-2 only showed minor heterogeneous distribution in a EC monolayer (Figure 3C). As ICAM-1 and VCAM-1 expression are both induced upon inflammation, we analysed whether ICAM-1 and VCAM-1 heterogeneity was correlated. Co-staining of ICAM-1 with either ICAM-2 or VCAM-1 showed no correlation between ICAM-1 and ICAM-2 (r = 0.044, p = 0.072) (Figure S2A). A weak positive correlation was found for ICAM-1 and VCAM-1 (r = 0.532, p < 0.001), even though we also observed significant populations of ICAM-1^high^/VCAM-1^low^ and ICAM-1^low^/VCAM-1^high^ ECs (Figure S2B). To study whether our findings were specific for TNF-α treatment, we treated ECs with other inflammatory mediators such as Lipopolysaccharide (LPS), Interferon (IFN)-γ and Interleukin (IL)-1β. The results revealed that under any inflammatory stimulus tested, ICAM-1 and VCAM-1 showed strong heterogeneous distribution whereas ICAM-2 only showed minor heterogeneous distribution (Figure S2C).

**Figure 3.**
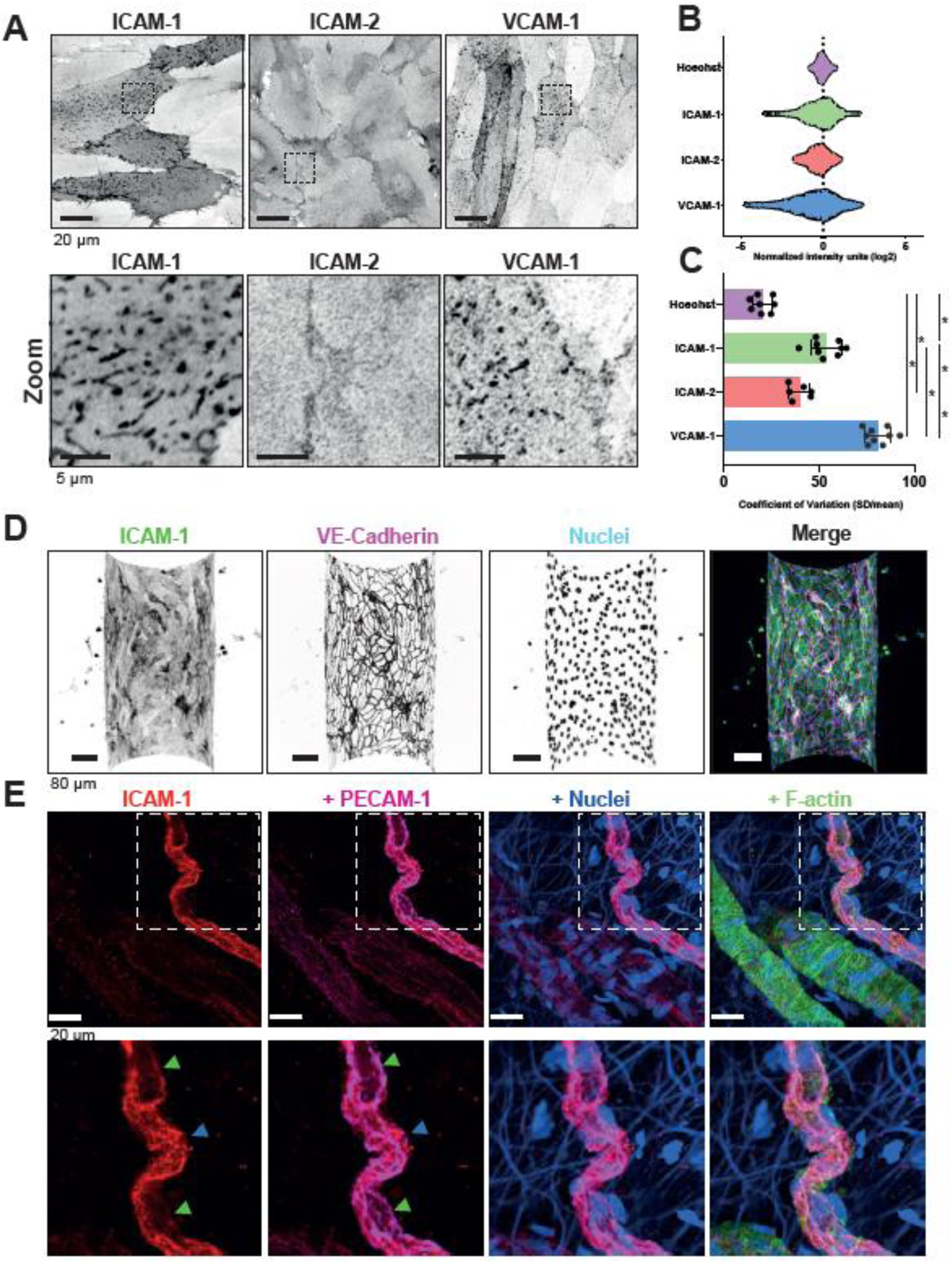
*Adhesion molecules display varying degrees of heterogeneity across varying conditions* **(A)** Inverted greyscale LUT of IF staining for ICAM-1, ICAM-2 and VCAM-1 on HUVECs after overnight TNF-α stimulation. ROIs represent zoom regions shown below. Scale bar, 20 µm in upper panels, 5 µm in bottom panels. **(B)** Violin plots showing Log2-normalized heterogeneous expression levels of Hoechst (N = 9 images, n = 1583 cells), ICAM-1 (N = 9 images, n = 1778 cells), ICAM-2 (N = 6 images, n = 1097) and VCAM-1 N = 9 images (n = 2306 cells). Each dot represents an individual cell and data is shown from 3 independent experiments. Data is normalized to mean intensity within an image to normalize for differences between each image. The dotted vertical line represents mean intensity **(C)** Bar graphs of the calculated coefficient of variation (CoV) (standard deviation/mean) for each field of view imaged in figure 2B. Data is shown from 3 independent experiments. Bar graphs show mean with SD. One-way ANOVA with multiple comparison correction was performed. ICAM-1 vs ICAM-2: p = 0.0018. All other combinations: p < 0.0001. **(D)** Inverted greyscale LUT of IF stain for ICAM-1, VE-cadherin and nuclei of a TNF-α treated vessel-on-a-chip composing of HUVECs. Scale bar, 80 µm. For clarity, only the bottom half of the Z-stack is shown. **(E)** *Ex vivo* whole-mount stains of colonic mesenterial adipose tissue of a patient with active inflammatory bowel disease, with ICAM-1 low (green arrow) and ICAM-1 high (blue) cells indicated. ICAM-1 is shown in red, PECAM-1 in magenta, nuclei in blue and F-actin in green. Scale bar, 20 µm.

To explore whether heterogeneous distribution of adhesion molecules in the inflamed EC monolayer changed over time, we allowed EC monolayers to mature for multiple days, ranging from 2 to 4 days, before treating with TNF-α. No change in the degree of heterogeneity of any of the adhesion molecules measured was found (Figure S2D). Using a vessel-on-a-chip model, developed by our lab^40^, we confirmed ICAM-1 heterogeneous distribution in a 3D inflamed vessel (Figure 3D and S2E). Clinically obtained samples of chronically inflamed human mesentery of inflammatory bowel disease patients showed ICAM-1 cell-to-cell heterogeneity in small veins (Figure 3E). Non-inflamed control tissue of the same organ, derived from intestinal carcinoma patients showed no ICAM-1 expression, but *ex vivo* treatment with TNF-α for 4h showed upregulation and heterogeneous distribution of ICAM-1 (Figure S2F). Together, these data show that heterogeneity of adhesion molecules is broadly conserved upon different conditions.

### ICAM-1 heterogeneity determines TEM hotspots

The functional existence of TEM hotspots is not clear. To understand this better, we focused on the major TEM hotspot marker ICAM-1 and generated stable ICAM-1 knockout (KO) ECs using Crispr/Cas9. In addition, we also generated ICAM-2 and ICAM-1/2 double knock out ECs. As HUVECs are limited by lifespan and passage time, we used blood outgrowth endothelial cells (BOECs) isolated from umbilical cord blood. These cells correspond to the characteristics ascribed to ECs and can be kept in culture for several passages^41^. Successful KO of ICAM-1 and -2 in ECs under TNF-α stimulation was confirmed by Western blotting (Figure S3A-C) and sequencing (Figure S3D). ICAM-1 KO did not influence ICAM-2 distribution compared to control ECs and ICAM-2 KO ECs still showed ICAM-1 heterogeneity (Figure S4A-B). Surprisingly, we found only a slight, non-significant decrease in neutrophil adhesion to ICAM-1-deficient ECs under flow conditions (Figure 4A). These data are in line with studies that used blocking antibodies against ICAM-1^42–44^. Depletion of ICAM-2 also did not alter neutrophil adhesion (Figure 4A). Interestingly, double KO ECs did show a 50% reduction of adhesion (Figure 4A). We did not find any effects on neutrophil diapedesis efficacy, as consistently around 80% of adhered neutrophils underwent diapedesis, indicating that ICAM-1 and/or -2 are not directly regulating neutrophil diapedesis (Figure 4B). Additionally, no effect on neutrophil crawling length, duration or speed was measured in any of the conditions (Figure S4C-E). Interestingly, when quantifying TEM hotspot events, we found a loss of TEM hotspots for neutrophils that adhered and crossed ICAM-1^-/-^ but not to ICAM-2^-/-^ EC monolayers (Figure 4C). This was found for the single as well as for the double KO conditions (Figure 4C).

**Figure 4.**
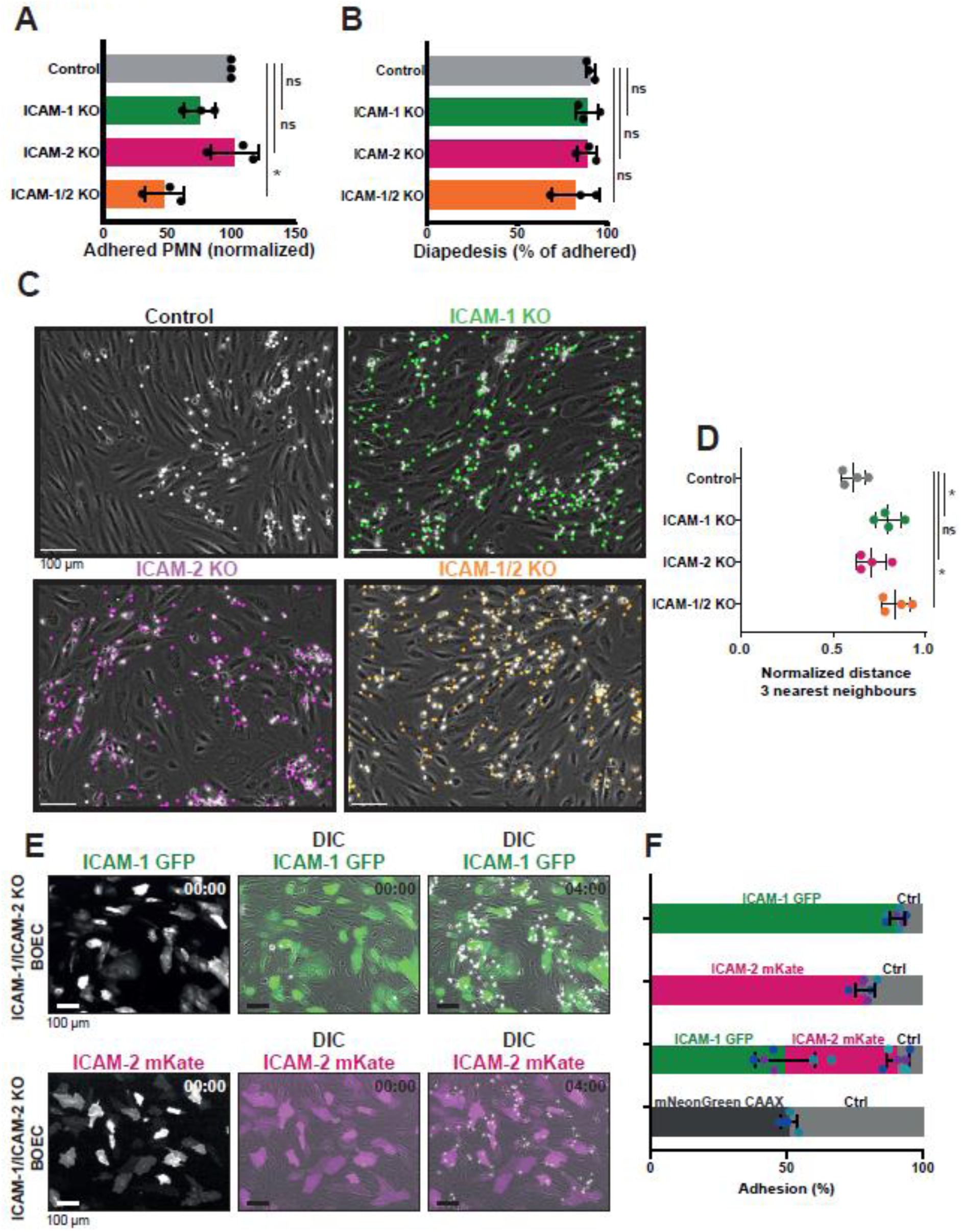
*ICAM-1 is the major marker for neutrophil hotspots.* **(A)** Quantification of number of adhered neutrophils (PMN) in TEM under flow assay using control BOECs (no gRNA), ICAM-1 KO BOECs, ICAM-2 KO BOECs, and double ICAM-1/2 KO BOECs. Data is normalized to control conditions (100%). Data consists of 3 independent experiments, 27141 total neutrophils measured. Bar graph displays mean with SD. One-way Paired ANOVA with multiple comparison correction, comparing all conditions with control. Control vs ICAM-1 KO: p = 0.6139. Control vs ICAM-2 KO: p = 0.9725. Control vs ICAM-1/2 KO: p = 0.0296. **(B)** Quantification of diapedesis efficacy (total transmigrated / total neutrophils detected *100%) of neutrophils through control, ICAM-1 KO, ICAM-2 KO, and ICAM-1/2 KO BOECs. Data consists of 3 independent experiments, 27141 total neutrophils measured. Bar graph displays mean with SD. One-way Paired ANOVA with multiple comparison correction, comparing all conditions with control. Control vs ICAM-1 KO: p = 0.9679 Control vs ICAM-2 KO: p = 0.9991. Control vs ICAM-1/2 KO: p = 0.2281. **(C)** Stills from neutrophil flow timelapses over control, ICAM-1 KO, ICAM-2 KO and ICAM-1/2 KO BOECs. All neutrophil TEM spots that occurred in the timelapse are shown in grey (control), green (ICAM-1 KO), magenta (ICAM-2 KO) and ICAM-1/2 KO (orange). Scale bar, 100 μm. **(D)** Medians of average distance of adhesion sites or TEM sites to 3 nearest neighbours, normalized against medians of the average distance to three nearest neighbours of the corresponding randomly generated spots. Data from 4 independent experiments is shown. One-way Paired ANOVA with multiple comparison correction, comparing all conditions with control. Control vs ICAM-1: p = 0.0150. Control vs ICAM-2: p = 0.2077. Control vs ICAM-1/2 KO: p = 0.0049. **(E)** Time lapse imaging of TEM under flow with ICAM-1/ICAM-2 KO cells. Part of EC monolayer is rescued with ICAM-1-GFP (green) or ICAM-2-mKate (magenta). Time indicated in the upper right corner in minutes. Left panels show ICAM-1 or ICAM-2 only channel, middle and right channel are merged with DIC. White dots are adhering neutrophils, predominantly at rescue cells. Scale bar, 100 μm. **(F)** Quantification of the preference for neutrophils to adhere to ICAM-1-GFP (green), ICAM-2-mKate (magenta) or CAAX-mNeonGreen (dark grey) expressed in ICAM-1/ICAM-2 KO ECs or ICAM-1/ICAM-2 KO ECs (Ctrl) (light grey). Bars represent percentage of neutrophil that adheres to indicated cell type. Numbers are corrected for area occupied. Dots are percentages from individual time lapse images, colours represent data from 3 independent experiments. Bars represent mean with standard deviation.

To confirm that heterogeneous distribution of ICAM-1 is crucial to induce TEM hotspots, we rescued the heterogeneous distribution of ICAM-1 in EC monolayers by overexpressing ICAM-1 in ICAM-1/ICAM-2 double KO-ECs in a mosaic fashion (Figure 4E and S4F). Interestingly, 90% of all neutrophils adhered to ICAM-1-GFP-expressing ECs but not to the KO-ECs (Figure 4F). Neutrophils also showed a preference for ICAM-2-expressing ECs, albeit less prominent compared to ICAM-1 (Figure 4F). Combining ICAM-1 and ICAM-2 heterogeneity showed that there was a small preference for ICAM-1 over ICAM-2 (Figure 4F). The membrane-marker CAAX was used as a control and did not affect the preference for adhesion (Figure 4F). These data indicate that ICAM-1 triggers TEM hotspots.

### TEM hotspots function to limit vascular leakage during TEM

One of the consequences of TEM hotspots is that less areas in the EC monolayer are penetrated by transmigrating neutrophils and consequently, the EC monolayer integrity can be maintained. Therefore, we hypothesized that TEM hotspots function to limit vascular leakage during TEM events. To test this, we measured permeability and neutrophil TEM simultaneously across ICAM-1/2 single and double KO-ECs using Transwell systems. No basal leakage was measured in any of the KO ECs when no neutrophils were present (Figure 5B and S5B). When measuring permeability during neutrophil TEM under control conditions, we did not find any change in permeability. However, we did measure an increase in EC permeability when neutrophils crossed EC monolayers that were deficient for ICAM-1 (Figure 5A-B). We also found an increase in permeability when neutrophils crossed the ICAM-1/2 KO EC monolayers, whereas permeability was only slightly increased when neutrophils crossed ICAM-2-deficient ECs (Figure 7A-B). TEM of neutrophils through these monolayers was consistent with TEM under flow experiments (Figure S5A and Figure 4A). These data indicate that TEM hotspots functionally protect endothelial monolayer integrity from leakage during TEM.

**Figure 5.**
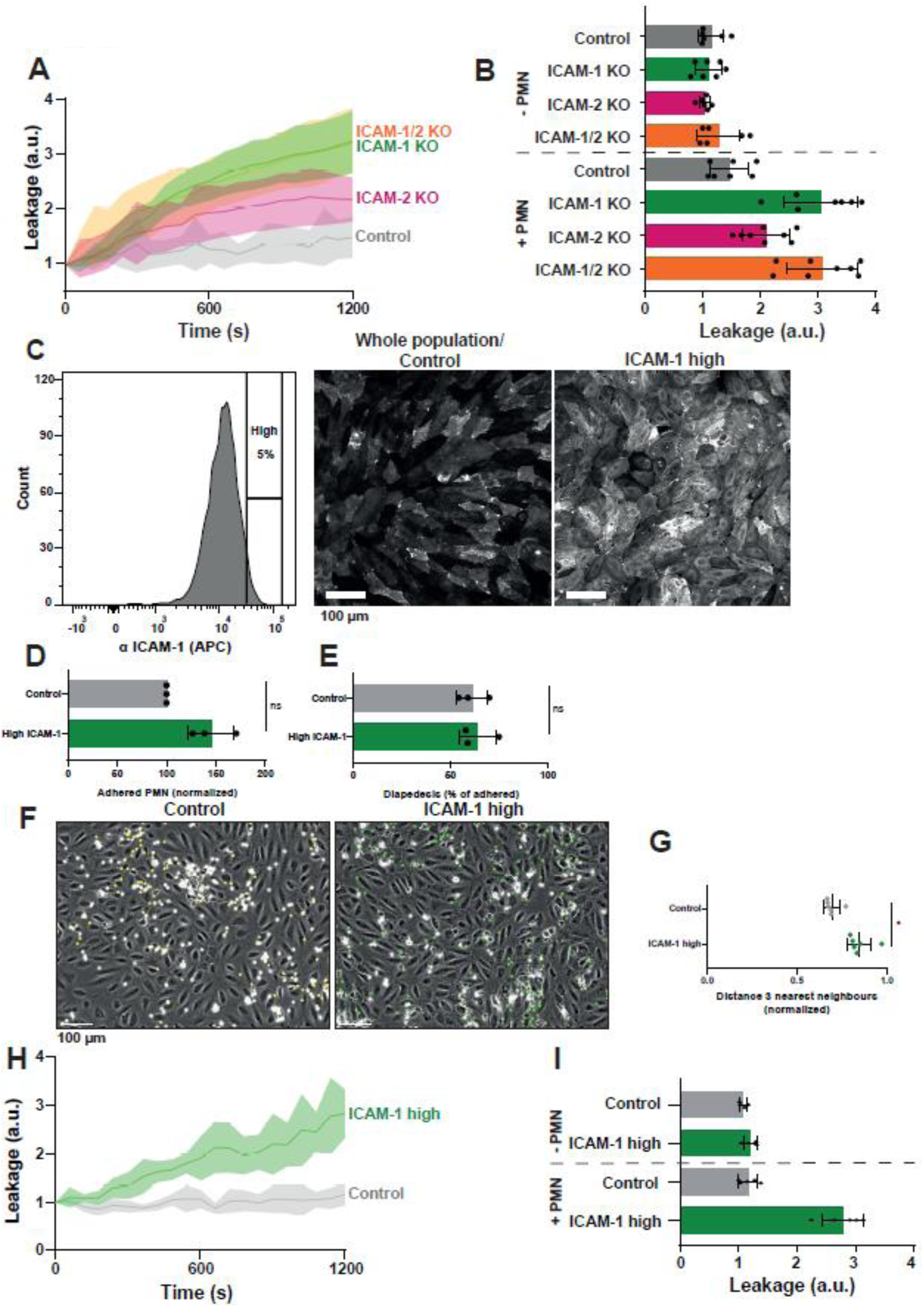
*ICAM-1 induced hotspots prevent vascular leakage.* **(A)** Texas-Red-dextran extravasation kinetics through control (grey), ICAM-1 KO (green), ICAM-2 KO (magenta) and ICAM-1/2 KO (orange) BOECs cultured on 3-μm pore permeable filters. DiO-stained neutrophils transmigrated towards C5a located in the lower compartment. Lines show means with 95% CIs of a total of 6 to 8 wells from 3 independent experiments. **(B)** Quantification of Texas-Red-dextran extravasation kinetics through control, ICAM-1 KO, ICAM-2 KO, and ICAM-1/2 KO BOECs after 20 minutes Bar graphs represent mean and SD. One-way ANOVA with multiple comparison corrections was performed within both the conditions without and with neutrophils. Without neutrophils (Control vs ICAM-1 KO: p = 0.9893, Control vs ICAM-2 KO: p = 0.8830, Control vs ICAM-1/2 KO: p = 0.7988, ICAM-1 KO vs ICAM-2 KO: p = 0.9689, ICAM-1 KO vs ICAM-1/2 KO: p = 0.5993, ICAM-2 KO vs ICAM-1/2 KO: p = 0.3768). With neutrophils (Control vs ICAM-1 KO: p < 0.0001, Control vs ICAM-2 KO: p = 0.1082, Control vs ICAM-1/2 KO: p < 0.0001, ICAM-1 KO vs ICAM-2 KO: p = 0.0068, ICAM-1 KO vs ICAM-1/2 KO: p = 0.9999, ICAM-2 KO vs ICAM-1/2 KO: p = 0.0042). **(C)** FACS graph displaying the top 5% sorted cells with the highest ICAM-1 expression levels based on fluorescence intensity. These cells are now called ICAM-1 high. **D)** Quantification of number of adhered neutrophils (PMN) in TEM under flow assay using control HUVECs and ICAM-1 high HUVECs. Data is normalized to control conditions (100%). Data consists of 3 independent experiments, 25646 total neutrophils measured. Bar graph displays mean with SD. Paired t-test: p = 0.0754. **(E)** Quantification of transmigration efficacy of neutrophils (PMN) (total transmigrated / total neutrophils detected *100%) through control HUVECs and ICAM-1 high HUVECs. Data is normalized to control conditions (100%). Data consists of 3 independent experiments, 25646 total neutrophils measured. Bar graph displays mean with SD. Paired t-test: p = 0.7923. **(F)** Stills from neutrophil flow timelapses over control, and ICAM-1 high HUVECs. All neutrophil TEM spots that occurred in the timelapse are shown in yellow (control) and green (ICAM-1 high). Scale bar, 100 μm. **(G)** Medians of average distance of adhesion sites or TEM sites to 3 nearest neighbours, normalized against medians of the average distance to three nearest neighbours of the corresponding randomly generated spots. Data of 6 videos from 3 independent experiments is shown. Mann-Whitney test: p = 0.0022. **(H)** Texas-Red-dextran extravasation kinetics through control (grey) and ICAM-1 high sorted (green) HUVECs cultured on 3-μm pore permeable filters DiO-stained neutrophils transmigrated towards C5a located in the lower compartment. Lines show means with 95% CIs of a total of 4 (without wells from 3 independent experiments. **(I)** Quantification of Texas-Red-dextran extravasation kinetics through control and ICAM-1 high sorted HUVECs. Bar graph displays mean with SD. Paired t-test was performed within conditions without and with neutrophils. Without neutrophils (Control vs ICAM-1 high: p = 0.1271). With neutrophils (Control vs ICAM-1 high: p < 0.0001).

To examine the effect of limiting leakage during TEM by reducing the heterogeneous expression of *endogenous* ICAM-1, we sorted the top 5% ICAM-1-expressing ECs, referred to as ICAM-1^high^ (Figure 5C). Indeed, the ICAM-1^high^ EC population displayed lower heterogeneity compared to control ECs (Figure S5C). As the antibody used for cell-sorting may interfere with neutrophil TEM, we tested its inhibitory properties. We observed no effect on adhesion, TEM and neutrophil crawling dynamics when the antibody was incubated for 24 hours, the same time the ECs are in contact with the antibody in the sorting experiments (Figure S5D-I). Using TEM flow assays, we observed that ICAM-1^high^ ECs showed increased, albeit non-significant, adhesion compared to control ECs (Figure 5D). Diapedesis efficacy was unaltered when comparing ICAM-1^high^ with control ECs (Figure 5E). However, when quantifying adhesion and diapedesis hotspots, we found that neutrophils showed less clustered transmigration patterns on ICAM-1-^high^ ECs compared to control ECs (Figure 5F). In line with these results, permeability assays demonstrated that EC leakage upon neutrophil TEM was increased in homogenous ICAM-1^high^-sorted ECs, in which no hotspots were detected (Figure 5H-I and Figure S5J-K). Thus, these data show that endogenous ICAM-1 heterogeneity is responsible for establishing functional TEM hotspots in the endothelium that limit vascular leak during TEM.

### The integrin binding domains of ICAM-1 are required for functional TEM hotspots

Neutrophils use integrins LFA-1 or Mac-1 to bind to ICAM-1 and ICAM-2, both using different epitopes. To study in more detail which of these epitopes are crucial for establishing functional TEM hotspots, we generated deletion mutants of ICAM-1, lacking one or more of the extracellular Ig-like domains. All deletion mutants for ICAM-1 were expressed in a heterogenous manner in ICAM-1 KO ECs and stained, on non-permeabilized samples, with an ICAM-1 antibody directed against the first Ig-like domain, showing normal distribution in apical filopodia (Figure S6A). To study which domains are crucial for TEM hotspot determination, we re-expressed these truncations in ICAM-1/2 KO-ECs and found that the lack of Ig-like domain 1 or 1-2 caused a mild decrease in TEM hotspots (Figure 6A). A similar decrease was observed when Ig-like domain 3 was depleted (Figure 6A). Interestingly, no TEM hotspot preference was measured when the first 3 Ig-like domains were deleted (Figure 6A). Ig-like domain 4, known for ICAM-1 dimerization^24^, had no effect on TEM hotspot preference (Figure 6A). Interestingly, deletion of the intracellular tail of ICAM-1, known to induce TEM-mediated signals^45–46^, did not influence hotspot recognition. These data showed that the first 3 Ig-like domains, the LFA-1 and Mac-1 epitopes of ICAM-1 are crucial for TEM hotspot determination and that ICAM-1 dimerization as well as intracellular signalling induced by the intracellular tail of ICAM-1 are not involved.

**Figure 6.**
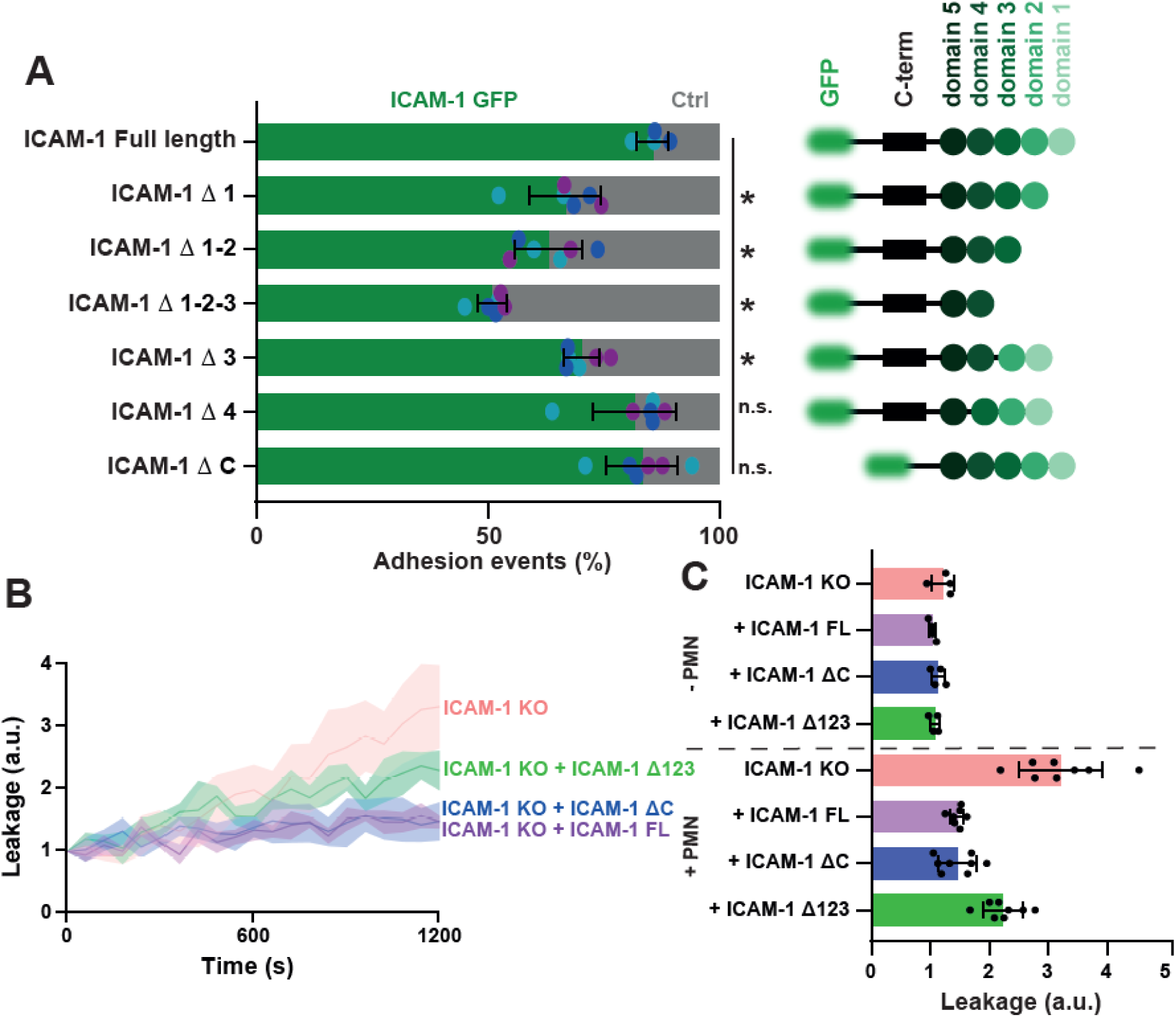
*Integrin-binding domains of ICAM-1 important for adhesion hotspots, the intracellular domain is not.* **(A)** Quantification of preference for neutrophils to adhere to ICAM-1-GFP truncations (green) expressed in ICAM-1/2 KO BOECs or ICAM-1/2 KO BOECs (Ctrl) (light grey). A schematic overview of all cloned ICAM-1 truncations is displayed as well. Bars represent the percentage of neutrophils that adhere to indicated transfected ECs. Numbers are corrected for area occupied. Dots are percentages from individual time lapse images. Bars represent mean with standard deviation. Colours represent data from 6 videos from 3 independent experiments (2 videos from 2 independent experiments for the control). One-way ANOVA with multiple comparison corrections, comparing all conditions to FL ICAM-1. FL vs Δ 1: p = 0.0004. FL vs Δ 12: p < 0.0001. FL vs Δ 123: p < 0.0001. FL vs Δ 3: p < 0.0029. FL vs Δ 4: p = 0.5848. FL vs Δ C: p = 0.6014. **(B)** Texas-Red-Dextran extravasation kinetics through ICAM-1/2 KO (pink) BOECs with mosaicly expressed ICAM-1-GFP (purple), ICAM-1-GFP Δ123 (green) and ICAM-1-GFP ΔC (blue), cultured on 3-μm pore filters. DiO-stained neutrophils transmigrated towards C5a located in lower compartment. Lines show means with 95% CIs of a total of 4 (without neutrophils) or 8 (with neutrophils) wells from 3 independent experiments. **(C)** Quantification of Texas-Red-dextran extravasation kinetics through ICAM-1/2 KO BOECs with mosaicly expressed ICAM-1-GFP, ICAM-1-GFP Δ123 and ICAM-1-GFP ΔC, after 20 minutes. Bar graphs represent mean and SD. One-way ANOVA with multiple comparison corrections was performed within both the conditions without and with neutrophils. Without neutrophils (ICAM-1 KO vs FL: p = 0.1845, ICAM-1 KO vs ΔC: p = 0.7368, ICAM-1 KO vs Δ123: p = 0.3441, FL vs ΔC: p = 0.6674, FL vs Δ123: p = 0.9717, ΔC vs Δ123: p = 0.8883). With neutrophils (ICAM-1 KO vs FL: p < 0.0001, ICAM-1 KO vs ΔC: p < 0.0001, ICAM-1 KO vs Δ123: p = 0.0006, FL vs ΔC: p > 0.9999, FL vs Δ123: p = 0.0057, ΔC vs Δ123: p = 0.0065).

To study whether the observed increase in permeability in previous experiments is due to the loss of ICAM-1-mediated TEM hotspots, we measured permeability during TEM across ICAM-1-deficient EC monolayers that were rescued with truncated mutants. ICAM-1-KO ECs that expressed the ICAM-1 mutant lacking the first 3 Ig-like domains (Δ123) showed a significant increase in leakage compared to ICAM-1-FL rescue conditions (Figure 6B-C). Total number of neutrophils that crossed the endothelial monolayers under these conditions was only marginally reduced (Figure S6B). Note that the ICAM-1 mutant that lacked the intra cellular tail did not show any increase in permeability. However, the number of neutrophils that crossed this monolayer were reduced, line with current literature (Figure S6B)^39^. None of the EC monolayers that expressed ICAM-1 mutants showed any basal leakage in the absence of neutrophils (Figure S6C).

Taken together, these data suggest that the binding of leukocytic integrins to endothelial ICAM-1 are required for TEM hotspot recogniti6r non. As a functional consequence, the existence of hotspots limits vascular permeability during TEM.

## Discussion

The existence of TEM hotspots has been recognized *in vivo*^11^, but there is no evidence for their biological relevance, nor is there clear consensus on the mechanism for hotspot recognition by leukocytes^12^. In this work, we use live-imaging and newly developed computational methods for the analysis of hotspots, providing new insight that addresses these questions. Our work confirms that TEM hotspots can also be found *in vitro*. As for the physiological relevance, we show for the first time that these hotspots on the endothelial monolayer limit vascular leakage during TEM. Mechanistically, this study shows that the first 3 extracellular Ig-domains of ICAM-1 are crucial for hotspot recognition.

Neutrophil TEM hotspots were first described *in vivo*, where the involvement of LFA-1 and Mac1 in different TEM phases was shown^11^. This study introduced the terms ‘Hotspot I’ for transendothelial migration hotspots and ‘Hotspot II’ for hotspots in the pericyte and basement membrane layer. In agreement with this study, we show that ICAM-1, ligand for LFA-1 and Mac-1, is involved in ‘Hotspot I’, resulting in local TEM.

Our major finding lies in the fact that TEM hotspots function to limit vascular leakage during TEM. The endothelium uses heterogeneous distribution of ICAM-1 to induce TEM hotspots for leukocytes. The basis of TEM hotspots is the initial adhesion of the leukocytes top the endothelium, particularly driven by ICAM-1. Depletion of ICAM-1 does not hamper efficient TEM but does increase vascular leakage during TEM. Previously, we showed that a F-actin-rich ring acts like an elastic strap around the perpetrating leukocyte to limit permeability during TEM^10^. This F-actin ring is under tension as it needs local activity of the small GTPase RhoA and downstream myosin activity. Our work furthermore indicated a role for ICAM-1 upstream from RhoA activation, albeit we were unable to directly link ICAM-1 function to the formation of the F-actin ring. We now find that ICAM-1 is crucial in the formation of TEM hotspots and thereby reduces local permeability. However, the ICAM-1 ΔC mutant shows that it is the disappearance of hotspots, and not the lack on downstream signalling towards the RhoA-mediated pore closure, that causes vascular leakage. These data indicate that the formation of the F-actin ring and the recognition of TEM hotspots through ICAM-1 distribution are uncoupled.

Other groups have identified signalling pathways in the endothelium that lead to the closure of the endothelial gap that is induced by the penetrating leukocyte. Braun and colleagues elegantly showed that platelet-derived Ang1 activates the endothelial Tie2 receptor, resulting in local activation of the FGD5-Cdc42 axis and closure of the gap^9^. Martinelli and colleagues have shown that local Rac1 activities are involved in the release of cellular tension signals that induce self-restorative ventral lamellipodia to heal barrier micro-wounds^47^. These mechanisms may all be triggered when leukocytes penetrate the endothelial monolayer^12, 48^. From a more efficient and energy saving cellular perspective, it makes sense to concentrate and minimize such signalling pathways to be able to keep the integrity of the vascular wall as good as possible.

We have compared neutrophil crawling dynamics around endothelial hotspots areas with non-hotspot areas. This revealed that neutrophils that used TEM hotspots showed much shorter crawling tracks. We hypothesize that these differences are due to higher ICAM-1 expression that capture neutrophils on the spot more efficiently. However, as we do not see extended crawling tracks on ICAM-1 KO ECs, it is likely that other factors are also involved, potentially regulated by leukocytes themselves. An interesting hypothesis to explore further involves the ability of neutrophils to leave behind ‘membrane trails’ that support migration of subsequent leukocytes^49^.

Surprisingly, our results show that in ICAM-1-depleted EC monolayers, the effect on total adhesion and transmigration is minimal. Only when both ICAM-1 and -2 are depleted, a clear decrease in neutrophil adhesion and therefore transmigration was observed, confirming earlier hypotheses that ICAM-1 and ICAM-2 have partly overlapping roles during TEM^50^. But importantly, our work adds new information to the separate roles of ICAM-1 and ICAM-2 in TEM. By overexpressing both adhesion molecules in a mosaic fashion, we show a preference for ICAM-1 over ICAM-2 for adhesion of neutrophils. This preference may be a result of the reported higher binding affinity of ICAM-1 with LFA-1^48^. Alternatively, ICAM-1 is enriched in apical filopodia that extend into the lumen and thus readily accessible for the rolling leukocyte, which is not the case for ICAM-2.

In this study, we highlight the significance of heterogeneous distribution of adhesion molecule protein expression within the endothelial monolayer for barrier integrity during TEM. Heterogeneous distribution of adhesion molecule such as ICAM-1 upon inflammation is broadly recognized and found *in vivo*^35^. Remarkably, based on our results, there does not seem to be a general correlation of heterogeneity between two inflammation-upregulated adhesion molecules ICAM-1 and VCAM-1, suggesting a more intricate mechanism that may work differently for each protein and each leukocyte subset. Earlier research has already provided clues at the epigenetic level: heterogeneity of well-known endothelial protein von Willebrand factor (VWF) is dependent on noise-induced changes in DNA methylation of the *VWF* promotor^51^, whereas VCAM-1 mosaic expression is due to heterogenous states of *VCAM-1* promotor methylation states^52^.

In conclusion, we have discovered how the endothelium takes advantage of adhesion molecule heterogeneous distribution within the endothelial monolayer by introducing TEM hotspots for leukocytes that function to limit vascular leakage during diapedesis and therefore maintain vascular integrity.

## Methods

### Plasmids

ICAM-1-GFP was described earlier^38^ and cloned into a lentiviral pLV backbone using SnaBI (ThermoFisher, FD0404) and XbaI (ThermoFisher, FD0684) / NheI (ThermoFisher, FD0973). To generate ICAM-1 truncated sequences, Gibson cloning (NEB) was performed on the pLV-ICAM-1-GFP plasmid. All constructs contain GFP as FP and the ICAM-1 signal peptide, consisting of amino acids Met1 to Ala27. ICAM-1 Δ 1 is truncated from Gln28 to Val109; ICAM-1 Δ 12 has a deletion from Gln28 to Phe212; ICAM-1 Δ 123 lacks Gln28 to Ile307; ICAM-1 Δ3 is truncated from Val213 until Ile307; ICAM-1 Δ 4 lacks Pro311 to Arg391; ICAM-1 Δ C terminates at Asn504. pLV-ICAM-2-mKate was constructed and packaged by VectorBuilder (Vector ID is VB200624-1164vtm). pLV-mNeonGreen-Caax and pLV-mScarletI-CAAX have been described earlier by us^18^. mEos4b-N1 was a gift from Micheal Davidson (Addgene # 54814; http://n2t.net/addgene:54814; RRID:Addgene_54814)^53^. mEos4b was PCR’ed out of the mEos4b-N1 vector, after which it was ligated into a pLV backbone using SnaBI and XbaI/NheI. All primers used in cloning are shown in Table 1.

**Table 1.**
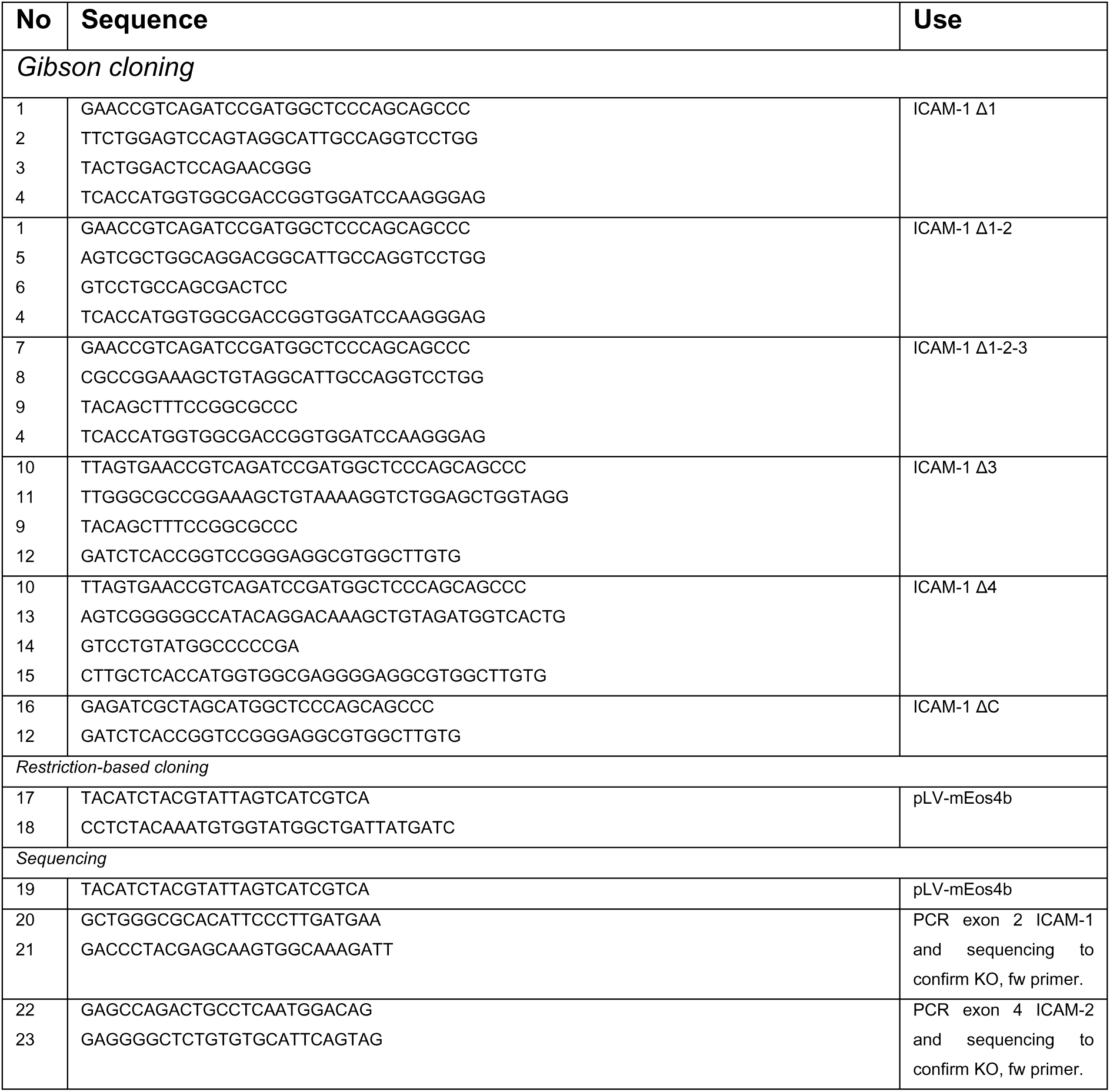
All primers used in this study.

### Antibodies

Alexa Fluor 647-conjugated ICAM-1 mouse monoclonal antibody was purchased from AbD Serotec (MCA1615A647T) (IF and FACS 1:400). Alexa Fluor 546-conjugated ICAM-1 mouse monoclonal antibody was bought from Santa Cruz (sc-107 AF546) (IF 1:400 Vessel-on-a-chip 1:200 whole-mount stain 1:100). FITC-conjugated ICAM-1 mouse monoclonal antibody was purchased from R&D (BBA20) (FACS 1:100). ICAM-1 rabbit polyclonal antibody was purchased from Santa Cruz (SC-7891) (WB 1:1000). ICAM-2 monoclonal mouse antibody was purchased from Invitrogen (14-1029-82) (IF 1:200). PE-conjugated ICAM-2 monoclonal mouse antibody was bought from BD (558080) (FACS 1:200). ICAM-2 rabbit monoclonal antibody was bought from Invitrogen (MA5029335) (WB 1:500). VCAM-1 monoclonal mouse antibody was purchased from Merck (MAB2511). Alexa Fluor 647-conjugated PECAM-1 monoclonal mouse antibody was bought from BD (561654) (whole-mount stain 1:200). Alexa Fluor 647-conjugated VE-cadherin mouse antibody was purchased from BD (561567) (Vessel-on-a-chip 1:200). Alexa Fluor 488-conjugated polyclonal chicken anti-mouse antibody (A21200) (IF 1:200) and Alexa Fluor 647-conjugated polyclonal chicken anti-mouse antibody (A21463) (IF 1:200) were purchased from Invitrogen. Alexa Fluor 488 phalloidin was purchased from Invitrogen (whole-mount stain 1:200). Hoechst 33342 (IF, vessel-on-a-chip and whole mount stain 1:50.000) was purchased from Molecular Probes (H-1399). Mouse monoclonal actin antibody for western blot (1:2500) was purchased from Sigma (A3853). Donkey anti-rabbit IRDye 800 (926-32213) (WB 1:5000) and donkey anti-mouse 680 (926-68022) (WB 1:5000) were purchased from LI-Cor. All antibodies were used according to manufacturer’s protocol.

### Cell culture and treatments

HUVEC were purchased from Lonza (C2519A) and cultured on fibronectin (FN)-coated dishes in Endothelial Growth Medium 2 (EGM-2) supplemented with singlequots (Promocell, C-22011) and 100 U/mL penicillin and streptomycin (P/S) at 37ᵒC in 5% CO_2_. HUVEC were cultured up to passage 7 and never allowed to grow above 70% confluency before the start of an experiment. Blood outgrowth endothelial cells (BOEC) were isolated from umbilical cord blood according to this protocol^41^. BOEC were grown on 0.1% gelatin-coated dishes during outgrowth and during experiments in EGM-2 supplemented with singlequots, 100 U ml^-1^ P/S and 18% fetal calf serum (Bodinco, Alkmaar, The Netherlands) at 37ᵒC in 5% CO_2_. HUVECs and BOECs were inflamed with 10 ng/mL recombinant TNF-α (Peprotech, 300-01A), 10 ng/mL IL-1β (Peprotech, 200-01B), 0.5 ng/mL IFN-γ (R&D, 285-IF-100) or 10 ng/mL LPS (Sigma, L2880) 20 hours before an experiment.

HEK-293T (ATCC) were cultured in Dulbecco’s Modified Eagle Medium (DMEM) (Gibco, 41965-039) containing 10% fetal calf serum, 100 U/mL P/S. By transfection of third generation lentiviral packaging plasmids with TransIT (Myrus, Madison, WI, USA) according to the manufacturers protocol, lentiviral particles containing pLV plasmids were generated. The second and third day after transfection, lentivirus-containing supernatant was harvested, filtered (0.45 micron) and concentrated with Lenti-X concentrator (Clontech, 631232). Virus was added to HUVEC or BOECs 1:250 to 1:500, depending on the efficacy of the virus. In case all cells were required to express the plasmid, a 2-day 1.5 μg/mL puromycin (InvivoGen, ant-pr-1) selection was performed. Endothelial cells were used in assays at least 72 hours after initial transduction.

### Generating ICAM knockout BOEC

ICAM-1 and ICAM-2 knockout BOEC were generated using guide RNAs (gRNAs) GCTATTCAAACTGCCCTGAT (ICAM-1) and GAGGTATTCGAGGTACACGTG (ICAM-2) that were ligated into a lentiviral Crispr vector (LentiCRISPRv2) containing Cas9 that was digested with BsmBI (ThermoFisher, FD0454) gRNA were designed using Crispr^54^. For ICAM-1, we targeted exon 2. For ICAM-2, we targeted exon 4. As a negative control, the CRISPR vector without a gRNA was used. Virus was produced and transductions in cord blood (BOEC were performed as described above. Transduced cells were selected using 1.5 μg/mL puromycin for 2 days, after which cells were single cell sorted into 96-well plates coated with 0.1% gelatin with an BD FACS Aria^TM^ III Cell Sorted (BD). For ICAM-2 knockout and for the ICAM-1/2 double knockout, ICAM-2 negative cells were sorted. For control gRNA, single cells positive for ICAM-2 were sorted. Since ICAM-1 only get expressed in inflammatory conditions, and BOECs stop growing after receiving inflammatory treatments, it was not possible to sort ICAM-1 knockout candidates with a fluorescent selection. Single cells were sorted and all monoclonal populations were tested for ICAM-1 expression when they reached around 100.000 cells, after which only the ICAM-1 lacking cell lines were kept in culture. Correct knockouts were, Western blot, FACS and genomic DNA extraction followed by sequencing using a DNeasy Blood & Tissue Kit (Qiagen, 69504) (Figure S3).

### Neutrophil isolation

Polymorphonuclear neutrophils were isolated from whole-blood, extracted from healthy voluntary donors that signed informed consent according to the rules maintained by the Sanquin Medical Ethical Committee, which are based on rules and legislation in place within The Netherlands. The rules and legislations were based on the Declaration of Helsinki (informed consent for participation of human subjects in medical and scientific research) and guidelines for Good Clinical Practice. Blood was always processed within 2 hours after donation. Whole blood is diluted 1:1 with 5% TNC in Phosphate Buffering Solution (PBS) (Fresenius Kabi, Zeist, The Netherlands) and pipetted on 12.5 mL Percoll (1.076 g/ml). Next, a 20-minute centrifugation (Rotina 420R) at 800xg with a slow start and no brake was performed on the diluted blood. After discarding the monocyte- and lymphocyte-containing ring fraction, 45 mL ice-cold erythrocyte lysis buffer (155 mM NH_4_CL, 10 mM KHCO_3_, 0.1 mM EDTA, pH7.4 in Milli-Q (Gibco, A1283-01)) was added to the pallet to lyse erythrocytes for 15 minutes. Erythrocyte lysis was performed twice, with a centrifuge step at 500xg for 5 min at 4ᵒC in between. Neutrophils were then centrifuged again at 500xg for 5 min at 4ᵒC, washed once with 30 mL ice-cold PBS, centrifuged again at 500xg for 5 min at 4ᵒC and resuspended in RT HEPES medium (20 mM HEPES, 132 mM NaCl, 6 mM KCL, 1 mM CaCl_2_, 1 mM MgSO_4_, 1.2 mM K_2_HPO_4_, 5 mM glucose (All Sigma-Aldrich), and 0.4% (w/v) human serum albumin (Sanquin Reagents), pH7.4). Neutrophil counts were determined using a cell counter (Casey). Neutrophils were kept at a concentration of 2 million/mL at RT. Neutrophils were kept no longer than 4 hours after isolation.

### Neutrophil transmigration under physiological flow

30.000 HUVECs or 20.000 BOECs per lane were seeded in respectively FN- or collagen-coated Ibidi μ-slides VI^0.4^ (Ibidi, Munich, Germany) and grown for 48 hours. TNF-α treatment (10 ng/mL) was performed 20 hours before the experiment, when the endothelial cells were grown into a confluent monolayer. A total of 6 million neutrophils, at 2 million/mL, was membrane-labelled for 20 minutes at 37ᵒC with Vybrant^TM^ DiO or DiD Cell-labeling solution (1:6000). Stained neutrophils were centrifuged for 3 minutes at 300xg at RT to wash away residual labeling solution. Neutrophils were resuspended in HEPES medium to a concentration of 1 million/mL. After letting the neutrophil recover at RT for 20 minutes, 1 million neutrophils at a time were incubated at 37ᵒC for 20 minutes before using them. The Ibidi flow chamber containing the endothelial cells was connected to a perfusion system and underwent shear flow of 0.5 mL/min (0.8 dyne/cm^2^) for 2 minutes before injecting 700.000 neutrophils into the tubing system.

Except for experiments with photoconvertible proteins, flow assays were imaged using an Axiovert 200 M widefield microscope, using a 10x NA 0.30 DIC Air objective (Zeiss). Fluorescent excitation light was provided by a HXP 120 C light source at 100% intensity and a TL Halogen Lamp at 6.06 V for transmitted light. Signal was detected with an AxioCam ICc 3 (Zeiss) camera. For the DIC channel, an exposure of 32 ms was used. For DiO-stained neutrophils, a 450-490 excitation filter, a 495 beam splitter, and a 500-550 emission filter were used with an exposure of 1900 ms. For DiD-stained neutrophils, a 625-655 excitation filter, a 660 beam splitter, and a 665-715 emission filter were used with an exposure of 1400 ms. To analyze neutrophil crawling dynamics and diapedesis locations Images were taken every 5 seconds for 15 minutes in two positions in the middle of the ibidi flow chamber lane. Immediately after acquiring the time-lapse, a tile scan of 4×6 frames was collected to quantify total adhesion and transmigration numbers. Images were taken using Zeiss using Zen Blue software. The tile scan was stitched using Zen Black software, using the DIC channel for stitching.

### Quantification neutrophil transmigration dynamics

All analyses were performed in Imaris version 9.7.2. To quantify total adhesion and diapedesis efficacy, a spot analysis was performed on the tile scans to count cells adhering on top of the monolayer and cells crawling on the subendothelial side of the monolayer. Spot analysis was performed on the DiD or DiO channel, with an estimated dot size of 8 micron. Only spots with a quality higher than 80 were filtered to ensure only neutrophils were counted. To distinguish neutrophils above and underneath the endothelium, a filter based on intensity in the DIC channel was added in the pipeline. Since neutrophils are white and round when on top of the endothelium and black and spread out when underneath the endothelium (Figure 1A), this filter could be used to separately count adhering and transmigrated neutrophils. Total adhesion was calculated as # adhering neutrophils + # transmigrated neutrophils, and due to large donor-dependent variation was normalized to a control experiment and shown as a percentage. Neutrophil diapedesis efficacy was quantified as (# adhering neutrophils/ (# total detected neutrophils) * 100%. The same spot analysis on neutrophils above the endothelial layer was performed on time-lapse data to quantify neutrophil crawling dynamics. A tracking step was added to the pipeline to connect the spots of each frame and detect neutrophil crawling patterns. For tracking analysis, the auto-regressive motion was used, with a maximum distance of 20 um between spots and allowing a gap size of 1 frame. Finally, tracks with less than 4 spots were filtered out to remove rolling neutrophils from the dataset. For optimal results, no more than 200 neutrophil tracks were allowed per video, and tracks were all manually checked for correctness. From this analysis, crawling speed, length, displacement, duration and linearity were calculated. Additionally, by taking the track starting location and track mean location, and subtracting those from each other, we were able to determine whether neutrophils crawled against or with the direction of flow. For this analysis, we discarded all tracks with less than 20 micron track lengths.

### Quantification of hotspot dynamics

Analysis of neutrophil crawling tracks in Imaris software was performed in widefield time-lapse data to compare behaviour of neutrophils at hotspots with neutrophils not utilizing hotspots. Only tracks that ended with diapedesis were used in this analysis. Tracks were classified as ‘hotspot tracks’ when they ended within 50 microns, the average diameter of a HUVEC cell, of another ending track. Neutrophils were classified as a ‘non-hotspot track’ if this criterium was not met. To assess the randomness of neutrophil diapedesis sites, track analysis was performed in Imaris on subendothelial neutrophils. All first spots of subendothelial crawling tracks were classified as ‘diapedesis sites’. All diapedesis site locations were masked in a new frame and a time projection was performed to generate one frame with all diapedesis sites. A spot analysis in Imaris was done on this frame to count the number of diapedesis sites in the time-lapse and the mean distance to its one, three, five or nine nearest neighbouring diapedesis sites was calculated. In FIJI (v1.52p)^55^, the same number of random spots was generated in an image with the same dimensions size as the time-cropped time-lapse frame. The same spot analysis and subsequent calculations were performed on this image. To create a parameter for randomness, the median distance to n nearest neighbours of diapedesis sites was divided by the median distance to n nearest neighbours of random sites. The more this value approaches 1, the more the diapedesis sites approach a purely random distributed pattern. The same analysis was done for measuring randomness of neutrophil adhesion sites, but instead of using the first spot of subendothelial tracks, the first spot of neutrophils crawling on top of the endothelium were used. Additionally, only tracks that ended in successful diapedesis were used in this analysis.

### Artificial hotspot neutrophil flow assay

To generate artificial adhesion molecule heterogeneity, ICAM-1/2 double knockout BOECs were transduced with (truncated) variants of ICAM-1 and ICAM-2. No puromycin selection was performed to preserve the non-transduced cells. Neutrophil flow assays were performed as described above, using unstained neutrophils. The DIC channel was imaged the same as described above. GFP was imaged with the same settings as DiO. mKate was imaged using a 559-585 excitation filter, a 590 beam splitter and a 600-690 emission filter, with an exposure time of 1200 ms. To quantify whether neutrophils preferred to adhere to transduced cells, neutrophils landing spots were manually analyzed, tallying whether a neutrophil that later underwent successful diapedesis adhered to a transduced or non-transduced cell. To take into account the variation in transduction efficacy between fields of view, the counted adhesion events were normalized against the percentage of the area in the field of view that was covered by transduced cells.

### Fixed immunofluorescent stains

For regular 2D cultered samples, HUVECs or BOECs were cultured in respectively FN- and collagen-coated Ibidi μ-slides VI^0.4^ (Ibidi, Munich, Germany). For fixation, 100 uL 4% Paraformaldehyde (PFA) in PBS++ was added to a drained flow chamber. Since all antibodies bound to extracellular epitopes, no permeabilization step was performed. Samples were blocked with 2% Bovine Serum Albumin (BSA) in PBS++. Primary antibodies were incubated for 1 hour at RT in PBS++, after which, if not working with directly conjugated antibodies, secondary antibodies were also incubated for 1 hour at RT. Between all fixation, blocking and staining steps, the flow chamber was washed three times with PBS++. If two primary antibodies were both raised in the same species, a three-step staining was performed: starting with an unconjugated primary antibody, followed by an accompanying secondary antibody, followed by second, directly conjugated antibody.

A Zeiss LSM 980 with Airyscan 2 module was used for detailed high-resolution confocal imaging of fixed samples, using a Plan-Apochromat 40x NA 1.3 oil DIC objective (Zeiss, 420762-9800-000) and a voxel size of 0.053 x 0.053 x 0.220 μm to capture Z-stacks. For all images, Multiplex SR-8Y settings were used and a GaAsP-PMT detector was used as a detector. GFP was excited using a 488 nm laser with a laser power of 0.2%, mKate was imaged using a 561 nm laser with 2.4% laser power, and Alexa Fluor 647 was excited with a 639 nm laser using 0.6% laser power. Images were acquired and 3D Airyscan-processed in Zen Blue version 3.3. Maximum projections were constructed in FIJI.

Vessel-on-a-chip samples were cultured and imaged as described here^40^. Patient tissue samples were received from the department of Pathology of the Amsterdam UMC, location AMC. All tissue samples were obtained with informed consent and according to Dutch guidelines for secondary used biological materials. Patient tissue samples were received from the department of Pathology of the Amsterdam UMC, location AMC. All samples were obtained and handled according current Dutch legislation regarding responsible secondary use of human tissues. Chronically inflamed patient samples, originating from the colonic mesentery, were obtained from inflammatory bowel disease patients, were obtained from inflammatory bowel disease patients during partial colectomy. Healthy mesenteric tissue was obtained from residual tissue of patients with intestinal carcinoma undergoing resection surgery. All tissue was stored in PBS++ at 4 degrees and prepared for imaging within 24 hours. Samples were prepared by cutting off small pieces of around 0.5 cm in diameter. If required, samples were incubated for 4 hours in PBS++ containing 10 ng/mL TNF-α for 4 hours at 37 °C. These pieces were fixed with 4% PFA for 15 min at 37 °C, permeabilized for 10 min with 0.5% triton-X at RT and finally blocked with 2% BSA for 30 min at RT. Between all steps, the sample was washed with PBS++. Stains were performed with a 1-hour incubation step by putting the sample in an antibody solution in a 1.5 mL Eppendorf in a slow rotator at 37 °C. Finally, tissue samples were mounted on glass bottom microwell dishes (MatTek, P35G-1.5-14-C) using 10% Mowiol.

Samples were imaged with the Zeiss LSM 980 with Airyscan module, using the same set-up as described above and having a voxel size of 0.038 x 0.038 x 0.170 μm. Hoechst was measured using a 405 nm laser with 2.4% laser power, ATTO-488 was measured with a 488 nm laser with 0.5% laser power, Alexa Fluor 568 was excited with a 561 nm laser with 4.5% laser power, and Alexa Fluor 647 was excited using a 639 nm laser with 8.5% laser power. All images were 3D Airyscan-processed.

### Confocal imaging of adhesion molecule heterogeneity

To image heterogeneity of adhesion molecules, fixed Ibidi flow chambers were stained for nuclei and ICAM-1, ICAM-2 or VCAM-1. Z-stack imaging was performed with the Zeiss LSM 980, using its confocal mode, using a Plan-Apochromat 20x objective NA 0.8 (Zeiss, 420650-9903-000). Voxel size was 1.184 x 1.184 x 0.5 μm. Hoechst was imaged with a 405 nm laser at 5.50% laser power and Alexa Fluor was excited with a 639 nm laser with 8% laser power. To measure heterogeneity, FIJI was used to generate sum projections of the Z-stacks. A rolling ball background subtraction of 25 pixels was performed on the nuclei channel, after which the nuclei were segmented by a threshold and a particle analysis on particles between 50-1000 pixels was performed. Then, fluorescent intensity was measured in the ICAM-1/ICAM-2/VCAM-1 channel. Data was normalized within every field of view to correct for inherent differences between fields of view. Coefficient of variation (SD/mean) of fluorescent intensity was used as a measurement of heterogeneity in the dataset.

### mEos4b photoconverting assay

For photoconversion assays, HUVECs were transduced and puromycin-selected with mEos4b. Neutrophil flow assays with unstained neutrophils were performed as described above. Imaging was performed on the Zeiss LSM 980, using its confocal mode as described above. First, neutrophil TEM was live-imaged for 15 minutes at three positions, imaging only the transmitted light channel. Afterwards, the same three positions were imaged fluorescently in channels designed to capture both the green and red emitting variant of mEos4b. For the green channel, a 488 nm laser with 4% laser power was used. For the red channel, a 561 nm laser with 5% laser power was used. Frames were captured every 8 seconds, and after the fourth frame the interactive bleaching module was utilized to photoconvert mEos4b in the whole field of view towards its red emitting variant. For photoconverting, the 405 nm laser was used at 50% for 5 seconds. After photoconversion, the Ibidi flow chambers were immediately fixed and stained as described above for nuclei and an adhesion molecule. To find back exactly the same fields of view that were live-imaged, we scanned the Ibidi flow chamber for photoconverted mEos4b and took images to measure heterogeneity of adhesion molecules. To correlate adhesion events to level of adhesion molecule expression, all ECs were numbered, after which the number of adhesion events followed by either diapedesis or detachment on each EC was tallied. Finally, all ECs on which adhesion events have taken place were ranked from 0% (lowest expression) to 100% (highest expression).

### Texas Red-Dextran permeability and transmigration assay

Endothelial cells (50.000 HUVEC or 35.000 BOEC) were seeded in FN-coated 24-well cell culture inserts (Corning FluoroBlok, Falcon, 3.0-μm pore size 351151) in a 24-well plate (Corning Companion Plate, Falcon, 353504) and cultured for 48 hours. Endothelial cells were treated with 10 ng/mL TNF-α 20 hours before the experiment. 100.000 DiO labeled neutrophils (1:6.000) and 100 μg Texas Red-Dextran (70 kDa; Sigma) in HEPES medium (20 mM HEPES, 132 mM NaCl, 6 mM KCL, 1 mM CaCl_2_, 1 mM MgSO_4_, 1.2 mM K_2_HPO_4_, 5 mM glucose (All Sigma-Aldrich), and 0.4% (w/v) human serum albumin (Sanquin Reagents, Amsterdam, The Netherlands), pH7.4) were added to the upper compartment of the culture insert in a total volume of 120 μL. 0.1 nM C5a (Sigma C-5788) in HEPES medium was added to the bottom compartment in a total volume of 600 μL. Leakage and neutrophil TEM were measured simultaneously for 20 minutes with an interval of 1 minute using an Infinite F200 pro plate reader (TECAN) at 37ᵒC. DiO labeled neutrophil TEM dynamics were measured using EX BP 490/9 and EM BP 535/20. Leakage dynamics of Texas Red-Dextran were measured with EX BP 595/9 and EM BP 630/20. To measure basal leakage, just Texas Red-dextran was added to the upper compartment.

### Western blotting

BOECs were grown in collagen-coated 6-well culture plates and washed twice with PBS++ (PBS containing 0.5 MgCl_2_ and 1 mM CaCl_2_). Lysis was performed with NP40 lysis buffer (50 mM TrisHCl, 100 mM NaCl, 10 mM MgCl_2_, 1% NP40 and 10% glycerol, pH7.4) with 1:500 protease inhibitor. Protein samples were centrifuged at 14.000 xG at RT for 10 minutes and resuspended in SDS-sample buffer containing 4% β-mecapto-ethanol. Samples were boiled at 95ᵒC for 3 minutes to denature proteins and separated on a 4-12% NuPage Bis-Tris gel (Invitrogen, NP0322BOX). Proteins were transferred using an iBlot Gel Transfer device (Invitrogen) for 7 minutes to a nitrocellulose membrane (Invitrogen, IB301002). Membranes were subsequently blocked with a 5% milk solution in tris-buffered saline with Tween 20 (TBST) at RT for 30 minutes. Primary antibodies were incubated overnight at 4 degrees in TBST and secondary IRDye 800 and IRDye 680 antibodies were incubated at RT for 1 hour. After each blocking and staining step, the membranes were washed with TBST 3x minutes. Western blots were developed using an Odyssey imaging system.

### FACS

FACS analysis was done using BD LSR II. Cells were detached using Accutase cell detachment solution (Sigma-Aldrich, A6964) and resuspended as 2 . 10^6^/ml in PBS containing 0.5% BSA, 1 mM CaCl_2_, 0.5 mM MgCl_2_. Antibody staining was performed for 30 minutes at 4°C. Cells were washed twice with PBS + BSA and directly analyzed. Single cells were gated using forward scatter and side scatter and further analyzed in FlowJo software (Tree Star, version 10).

### Cell sorting ICAM-1 high cells

HUVECs were grown into confluent monolayers in a total of 3 t150 flasks. After two washing steps with PBS++, HUVECs were stained for ICAM-1. Antibody stains for ICAM-1 were performed before cell detachment to mimic the ICAM-1 heterogeneity observed in monolayers most optimally. ICAM-1 antibody was incubated with the monolater for 20 min in EGM-2 at 37 °C. After two more washing steps with PBS++, 5 mL accutase was added to each t150 for 7 min at RT to take cells into suspension. Next, 5 mL EGM-2 was added to the suspension and cells were spinned down at 200xg for 5 min. The cell pallets were resuspended in EGM-2 in polypropylene tubes. A FACSAria^TM^ II or III Cell Sorter (BD) was used to sort the ICAM-1 5% high cells into suspension. As a control, the whole living cell population was sorted. After sorting, cells were seeded into Ibidi flow chambers or on transwell inserts, and neutrophil flow experiments and dextran leakage assays were conducted as described above.

### Statistics

Data are presented as either means or medians + SD, indicated for each graph. For neutrophil quantifications, comparisons between two groups were performed by a paired t-test and comparisons between multiple groups were performed by One-way paired ANOVAs, pairing data of a single donor. For other experiments, a student t-test or One-way ANOVA was performed, indicating which conditions were compared. For calculation correlations, Pearson r was calculated. A two-tailed P value of <0.05 was considered significant. For microscopy images, representative images are shown.

## Abbreviations

BOEC: Blood Outgrowth Endothelial Cells
BSA: Bovine Serum Albumin
DMEM: Dulbecco’s Modified Eagle Medium
EGM: Endothelial Growth Medium
FN: Fibronectin
gRNA: Guide RNA
HUVEC: Human Umbilical Vein Endothelial Cells
ICAM-1/2: Intercellular Adhesion Molecule
Ig: Immunoglobulin
IFN: Interferon
IL: Interleukin
LFA-1: Lymphocyte-Associated Antigen 1
LPS: Lipopolysaccharide
Mac-1: Macrophage-1
Antigen PBS: Phosphate Buffering Solution
P/S: Penicillin/Streptomycin
SDM: Site-Directed Mutagenesis
TBST: Tris-buffered saline with Tween 20
TEM: Transendothelial Migration
TNF: Tumor Necrosis Factor
VCAM-1: Vascular Cell Adhesion Molecule 1
VWF: von Willebrand Factor

## Conflicts of Interest

The authors declare no conflict of interest.

## Acknowledgements

This work was supported ZonMW NWO Vici grant # 91819632 (JDvB & MLBG)

---

**Figure S1.**
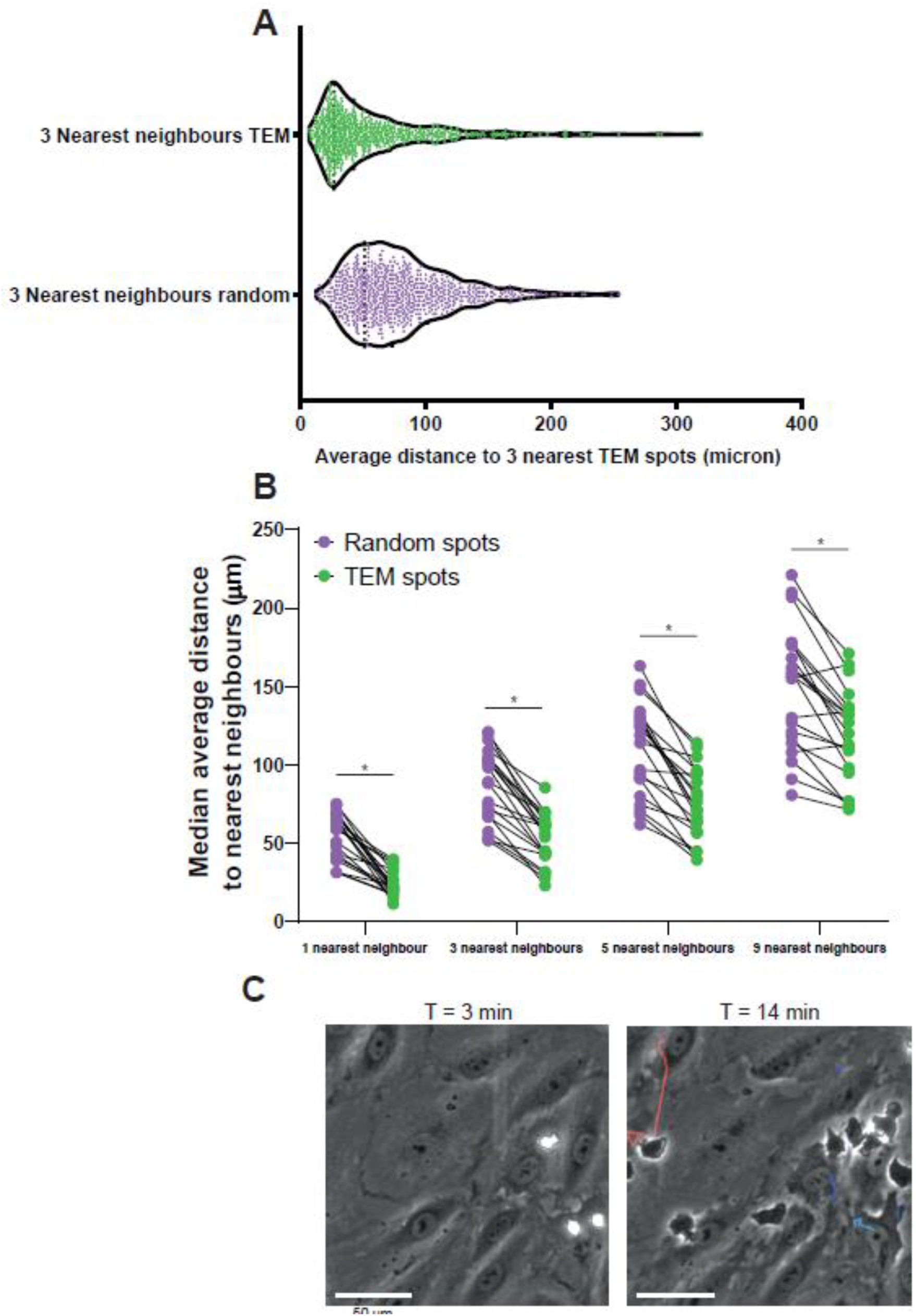
*Comparison of different nearest neighbour numbers and example hotspot and non-hotspot tracks*. **(A)** Violin plot of average distance to 3 nearest neighbours for actual diapedesis sites and randomly generated spots. Each datapoint corresponds to 1 diapedesis site and 729 datapoints from 21 time-lapses are plotted from 3 independent experiments. **(B)** Comparison of analysis methods for nearest neighbour calculations. 1, 3, 5 and 9 nearest neighbour(s) for each TEM spot were calculated and compared to randomly generated spots. Data is from 21 videos from 3 independent experiments. Paired t-test on the 21 medians for every comparison: p < 0.0001 for all. **(C)** Stills from a DIC time-lapse TEM assay, showing neutrophils and their complete crawling tracks at hotspots, indicated with blue tracks, and non-hotspots, indicated with a red track. The direction of flow is from top to bottom, time is indicated in min:sec at the top right. Scale bar, 50 µm.

**Figure S2.**
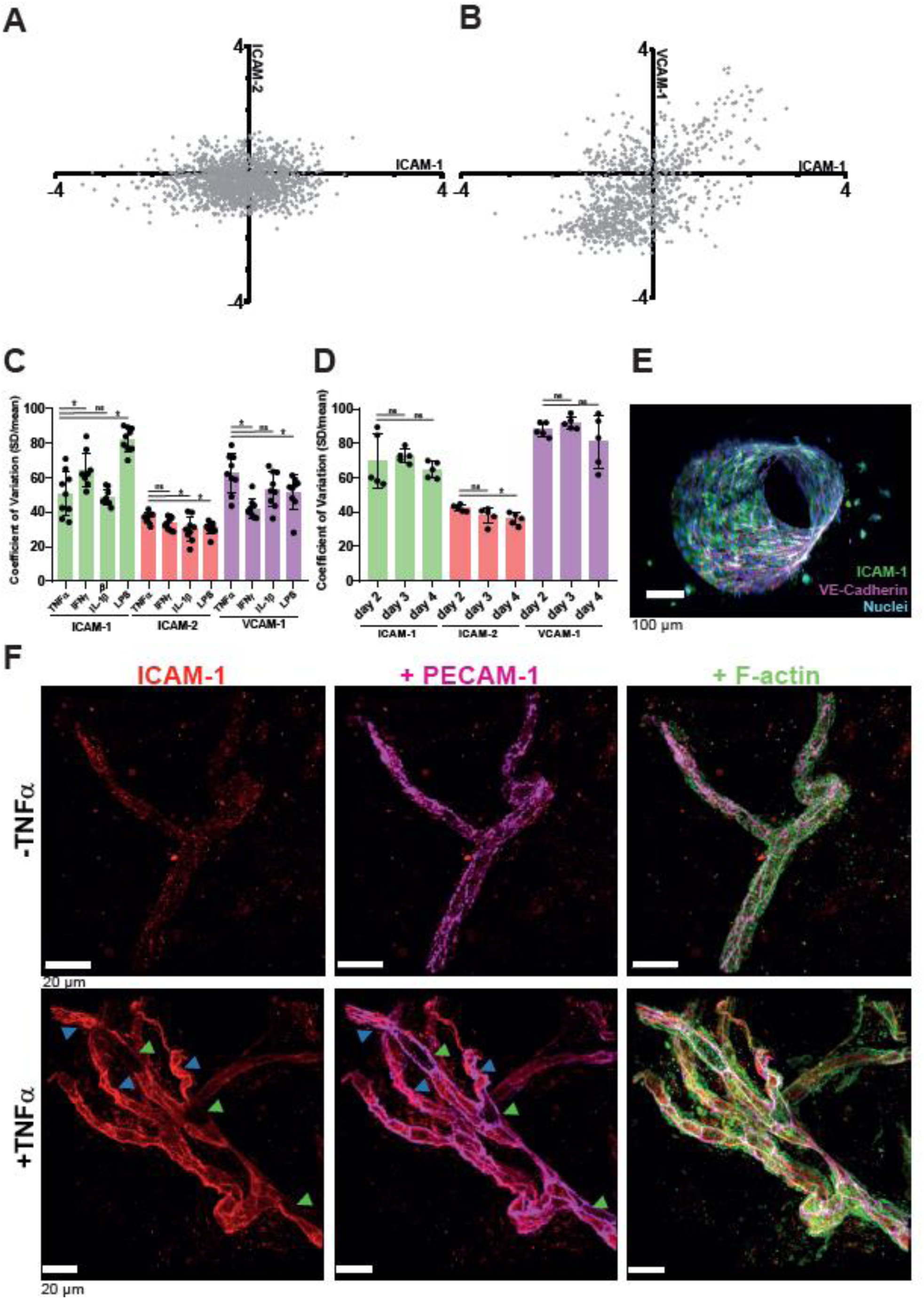
*Adhesion molecule heterogeneity persist across different variables and does not correlate strongly with each other.* **(A)** Correlation plot of Log2-transformed fluorescent intensity of ICAM-1 and ICAM-2, normalized within each field of view. r = 0.04481, p = 0.072 **(B)** Correlation plot of Log2-transformed fluorescent intensity of ICAM-1 and VCAM-2, normalized within each field of view. r = 0.532, p < 0.001 **(C,D)** Bar graphs displaying Coefficient of Variations (CoV) of ICAM-1, ICAM-2 and VCAM-1 for each field of view measured. Data is shown from 3 independent experiments. Bars show mean with SD. **(C)** Different inflammatory stimulants. One-way ANOVA on means with multiple comparison correction against TNF-α data, separate test for each protein (N = 9 images per condition). ICAM-1 (TNF-α vs IFN-γ: p = 0.0102. TNF-α vs IL-1β: p = 0.9171. TNF-α vs LPS: p < 0.0001). ICAM-2 (TNF-α vs IFN-γ: p = 0.2900. TNF-α vs IL-1β: p = 0.0107. TNF-α vs LPS: p = 0.0224). VCAM-1 (TNF-α vs IFN-γ: p = 0.0002. TNF-α vs IL-1β: p = 0.1192. TNF-α vs LPS: p = 0.0499). **(D)** Different maturation states of the endothelial monolayer. One-way ANOVA on means with multiple comparison correction against day 2 data, separate test for each protein (N = 5 images per condition). ICAM-1 (day 2 vs day 3: p = 0.8627. day 2 vs day 4: p = 0.6332.) ICAM-2 (day 2 vs day 3: p = 0.1338. day 2 vs day 4: p = 0.0262.) VCAM-1 (day 2 vs day 3: p = 0.7364. day 2 vs day 4: p = 0.4043.) **(E)** Side view of whole vessel-on-a-chip, shown in figure 2D. Stained for ICAM-1 (green), VE-cadherin (magenta) and nuclei (blue). Scale bar, 20 µm**. (F)** *Ex vivo* whole-mount stains of healthy mesenterial adipose tissue of a carcinoma patient, incubated with without or with 10 ng/µL TNF-α for 4 hours. ICAM-1 low (green arrow) and ICAM-1 high (blue) cells indicated. ICAM-1 is shown in red, PECAM-1 in magenta and F-actin in green. Scale bar, 20 µm.

**Figure S3.**
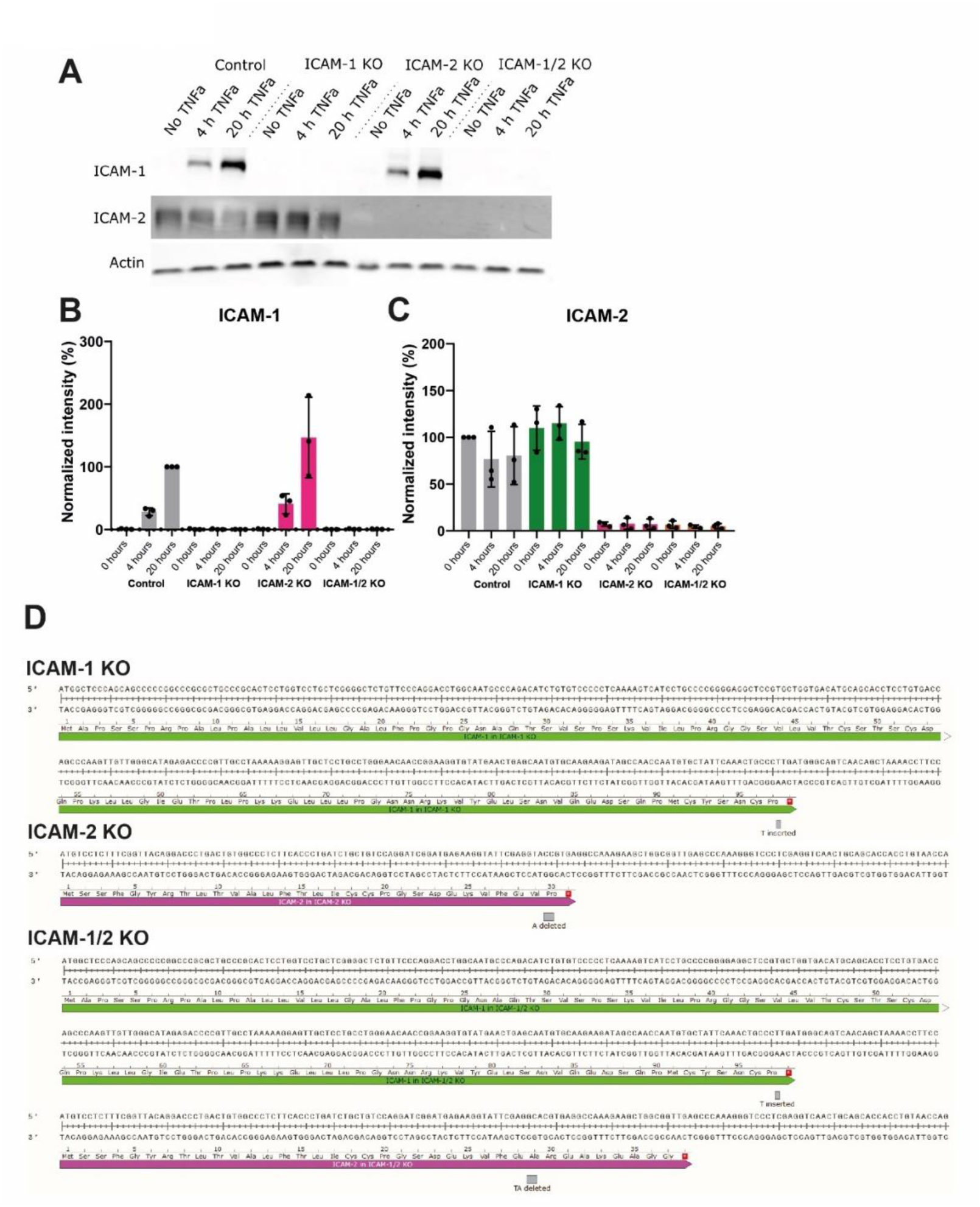
*Verification of ICAM-1/2 knockout BOECs.* **(A)** Western blot for ICAM-1 and ICAM-2 in Crispr knockouts after 0, 4 and 20 hours TNF-α. One representative gel out of three performed gels is shown. Actin is used as loading control. **(B,C)** Quantification of western blot for ICAM-1 **(B)** and ICAM-2 **(C)**, normalized to the actin loading control. Bar graphs display mean with SD. ICAM-1 data is normalized to 20 hours TNF-α in control BOECs, ICAM-2 data is normalized to 0 hours TNF-α in control BOECs. **(D)** Sequencing results of knockout cells. Mutations are depicted in grey block. Premature stop codon is depicted as star in red box. The protein sequences of the truncated ICAM-1 and ICAM-2 that are brought to expression in the KO BOECs are shown above the nucleotide sequence.

**Figure S4.**
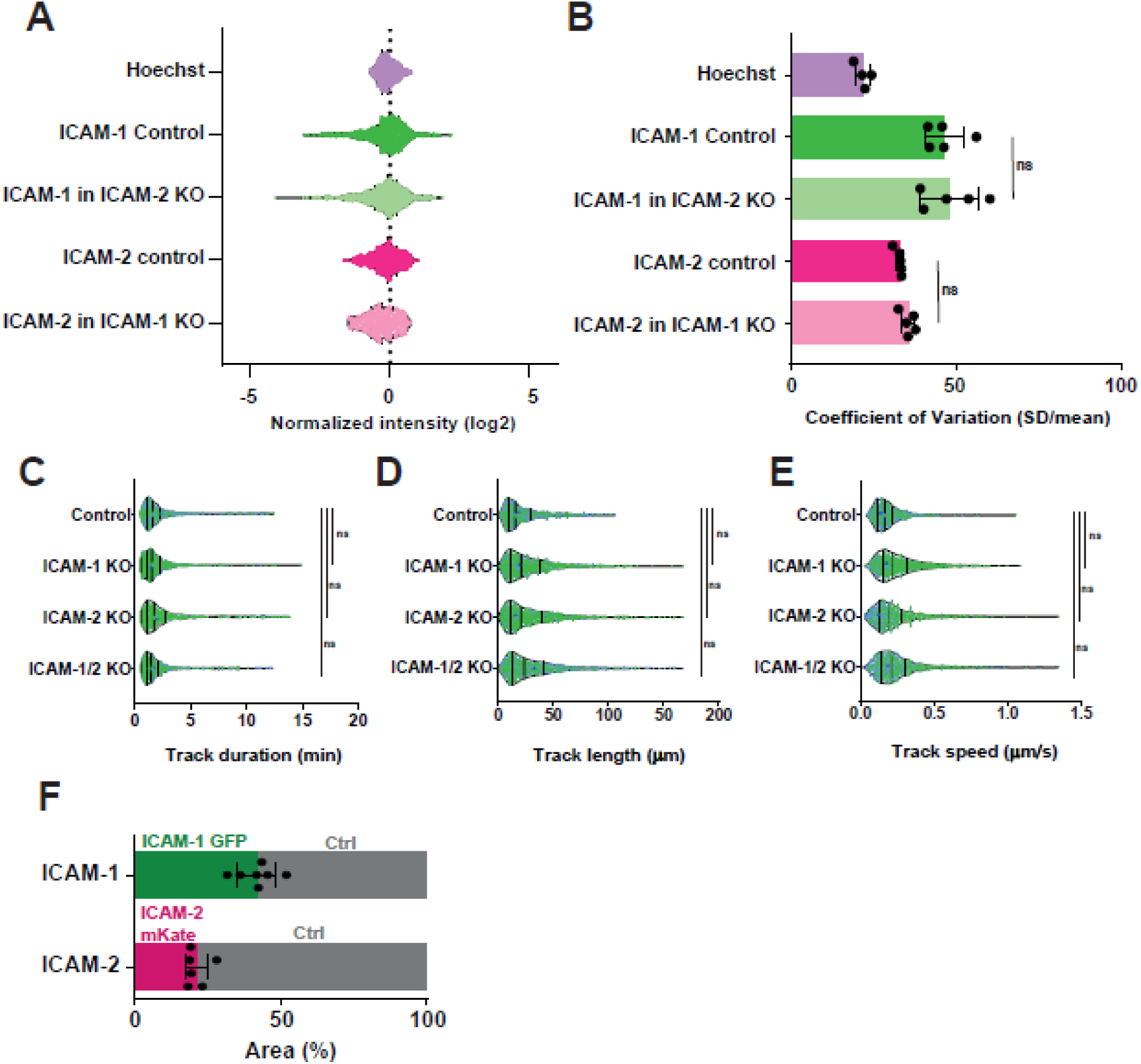
*ICAM-1 and ICAM-2 knockout does not influence each other’s heterogeneity and ICAM-1/2 KO BOECs do not have altered crawling dynamics.* **(A)** Violin plots displaying Log2- normalized fluorescent intensity of Hoechst in control (n = 488 cells), ICAM-1 in control (n = 611 cells) and in ICAM-2 KO BOECs (n = 497 cells), and ICAM-2 in control (n = 808 cells) and in ICAM-1 KO BOECs (n = 336 cells). Data is from 2 independent experiments **(B)** Bar graph displaying coefficients of variation (CoV) of all field of views measured in figure S4A. Mann-Whitney test between ICAM-1 stain conditions (p > 0.9999) and ICAM-2 stain condition (p = 0.0952). **(C,D,E)** Violin plots displaying neutrophil crawling duration, length and speed across control (n = 834 neutrophils), ICAM-1 KO (n = 1096 neutrophils), ICAM-2 KO (n = 1036 neutrophils) and ICAM-1/2 KO (n = 793 neutrophils) BOECs. Colours represent data from three independent experiments. Medians of three individual experiments are shown in bigger dots. Medians and quartiles of all data is displayed with vertical lines. One-way Paired ANOVA with multiple comparison correction, comparing all conditions with control. **(C)** Control vs ICAM-1: p = 0.3906. Control vs ICAM-2: p = 0.9299. Control vs ICAM-1/2 KO: p = 0.4066 **(D)** Control vs ICAM-1: p = 0.6582. Control vs ICAM-2: p = 0.9073. Control vs ICAM-1/2 KO: p = 0.2270 **(E)** Control vs ICAM-1: p = 0.0779. Control vs ICAM-2: p = 0.7565. Control vs ICAM- 1/2 KO: p = 0.0603 **(F)** Quantification of area of heterogeneous EC monolayer composing of ICAM-1/ICAM-2 KO ECs either non-expressing (ctrl, grey) or expressing ICAM-1-GFP (upper, green) or ICAM-2-mKate (lower, magenta).

**Figure S5.**
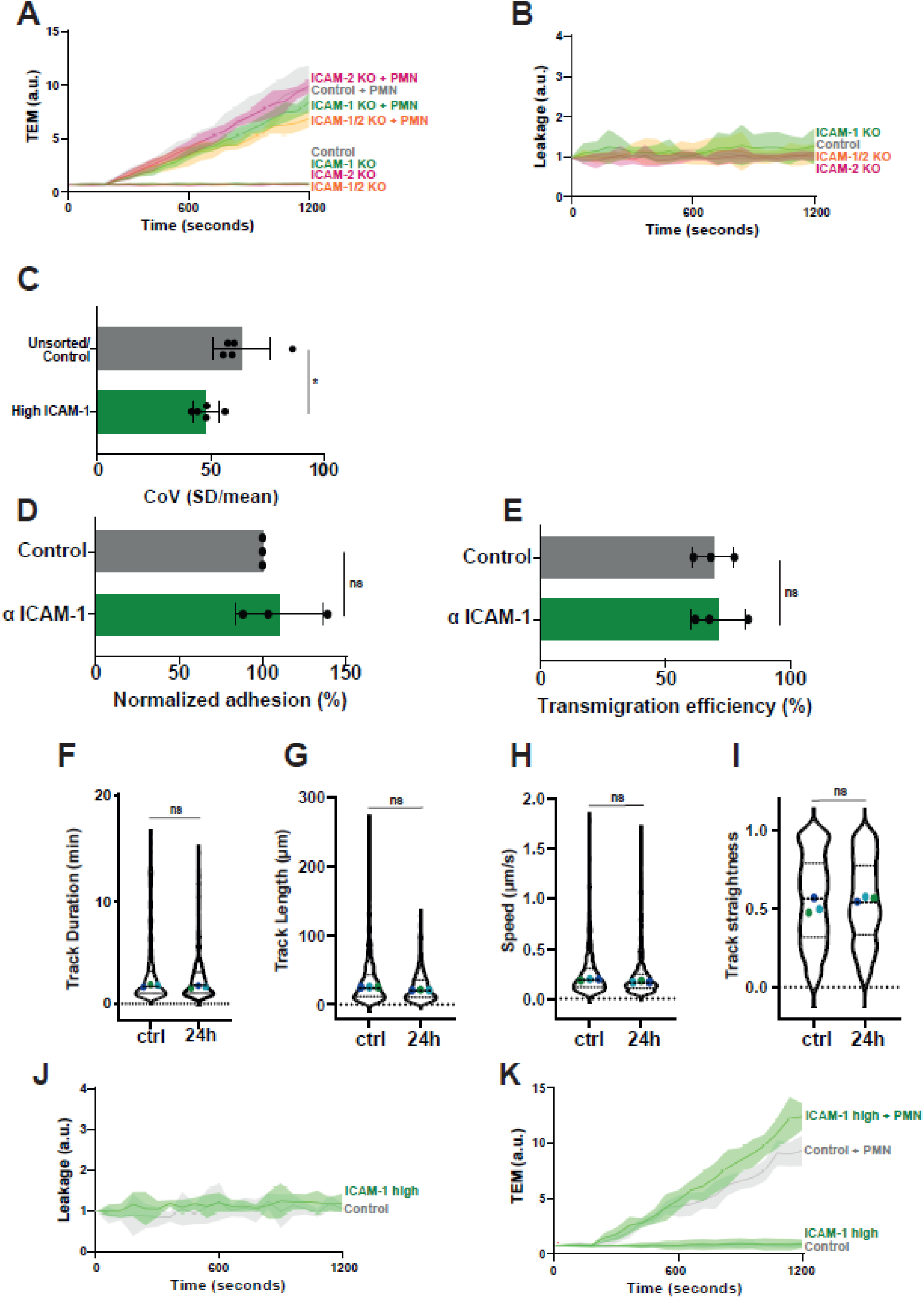
*Antibody against ICAM-1 used for sorting does not affect neutrophil TEM dynamics.* **(A)** Neutrophil extravasation kinetics through control (grey), ICAM-1 KO (green), ICAM-2 KO (magenta) and ICAM-1/2 KO (orange) BOECs cultured on 3-μm pore permeable filters. DiO-stained neutrophils transmigrated towards C5a located in the lower compartment. Lines show means with 95% CIs of a total of 6 to 8 wells from 3 independent experiments. **(B)** Basal leakage measured with Texas-Red-dextran extravasation kinetics through control (grey), ICAM-1 KO (green), ICAM-2 KO (magenta) and ICAM-1/2 KO (orange) BOECs. Lines show means with 95% CIs of a total of 6 to 8 wells from 3 independent experiments. **(C)** Bar graphs of Coefficient of Variation (CoV) of unsorted control and ICAM-1 high HUVECs. Mann-Whitney test: p = 0.0159. **(D)** Quantification of number of adhered neutrophils (PMN) in TEM under flow assay using control HUVECs and HUVECs incubated for 24 hours with an αICAM-1 antibody. Data is normalized to control conditions (100%). Data consists of 3 independent experiments, 7038 total neutrophils measured. Bar graph displays mean with SD. Paired t-test: p = 0.5646. **(E)** Quantification of transmigration efficacy of neutrophils (PMN) (total transmigrated / total neutrophils detected *100%) through control HUVECs and HUVECs incubated for 24 hours with an αICAM-1 antibody. Data is normalized to control conditions (100%). Data consists of 3 independent experiments, 7038 total neutrophils measured. Bar graph displays mean with SD. Paired t-test: p = 0.6801. **(F,G,H,I)** Violin plots displaying track duration, length, speed and straightness of neutrophils crawling on control HUVECs (629 neutrophils measured) and HUVECs (486 neutrophils measured) incubated for 24 hours with an αICAM-1 antibody. Medians and quartiles are shown, and three dots are medians of each independent experiment. Paired t-test on the medians. **(F)** p = 0.4094 **(G)** p = 0.2226 **(H)** p = 0.2736 **(I)** p = 0.9957 **(J)** Neutrophil extravasation kinetics through control (grey) and ICAM-1 high sorted (green) HUVECs cultured on 3-μm pore permeable filters. DiO-stained neutrophils transmigrated towards C5a located in the lower compartment. Lines show means with 95% CIs of 3 wells from 3 independent experiments. **(K)** Basal leakage measured with Texas-Red-dextran extravasation kinetics through control (grey) and ICAM-1 high sorted (green) HUVECs BOECs. Lines show means with 95% CIs of 3 wells from 3 independent experiments.

**Figure S6.**
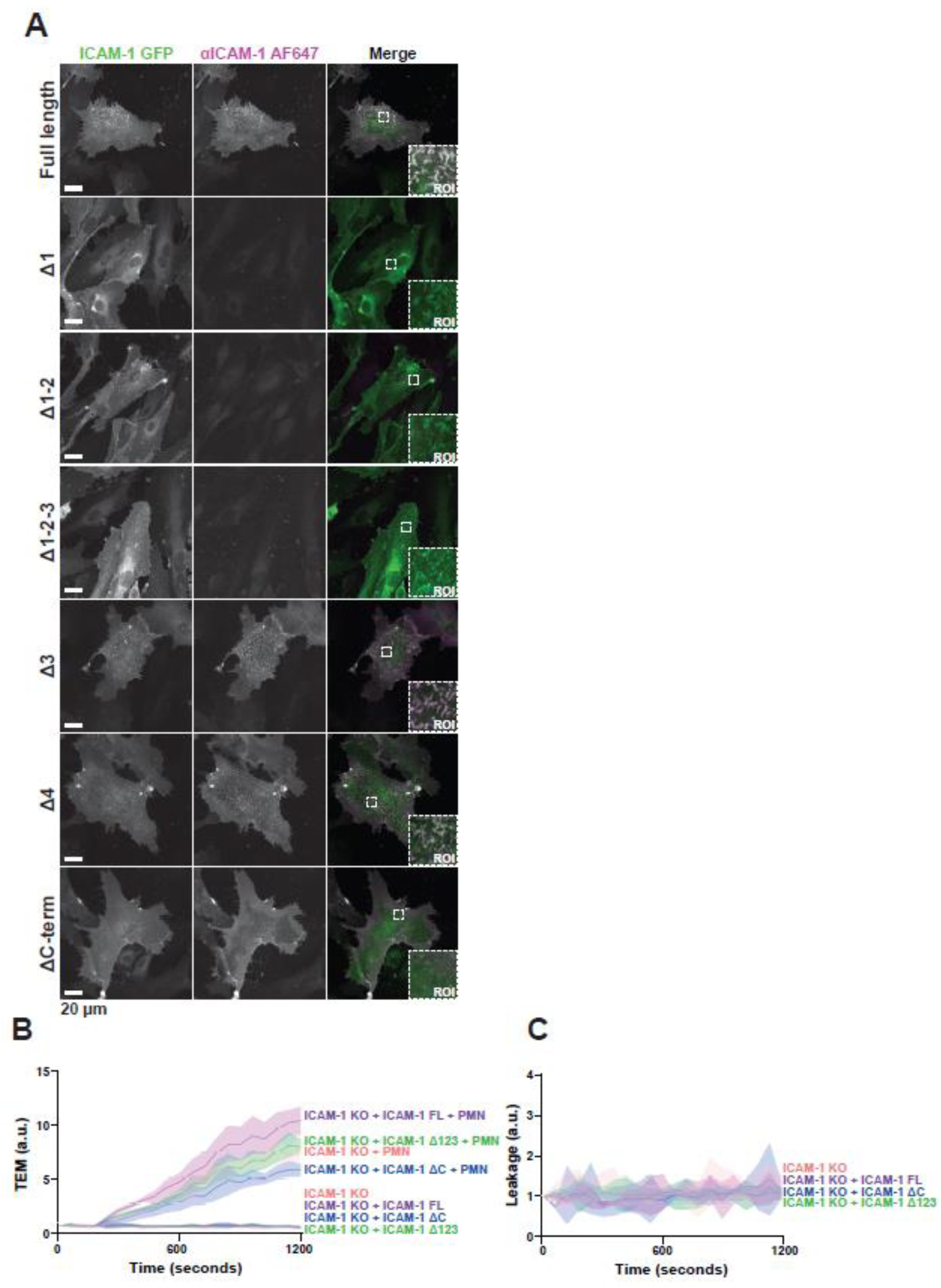
*ICAM-1 truncations localization in ECs*. **(A)** Immunofluorescence staining for ICAM-1 Ig-like extracellular domain 1 on all ICAM-1-GFP truncated proteins. Left panel shows ICAM-1-GFP truncated proteins (green), middle pannel shows IF staining for ICAM-1 Ig-like domain 1 (magenta) and right pannel is composite image of both. **(B)** Neutrophil extravasation kinetics through ICAM-1/2 KO (pink) BOECs with mosaicly expressed ICAM-1-GFP (purple), ICAM-1-GFP Δ123 (green) and ICAM-1-GFP ΔC (blue), cultured on 3-μm pore permeable filters. DiO-stained neutrophils transmigrated towards C5a located in the lower compartment. Lines show means with 95% CIs of a total of 4 (without neutrophils) or 8 (with neutrophils) wells from 3 independent experiments. **(C)** Basal leakage measured with Texas-Red-dextran extravasation kinetics through ICAM-1/2 KO (pink) BOECs with mosaicly expressed ICAM-1-GFP (purple), ICAM-1-GFP Δ123 (green) and ICAM-1-GFP ΔC (blue). Lines show means with 95% CIs of 4 wells from 3 independent experiments.

